# The genetic architecture of adaptation to leaf and root bacterial microbiota in *Arabidopsis thaliana*

**DOI:** 10.1101/2022.09.26.509609

**Authors:** Fabrice Roux, Léa Frachon, Claudia Bartoli

## Abstract

Understanding the role of host genome in modulating microbiota variation is a need to shed light into the holobiont theory and overcome the current limits on the description of host-microbiota interactions at the genomic and molecular levels. However, the host genetic architecture structuring microbiota is only partly described in plants. In addition, most association genetic studies on microbiota are often carried out outside the native habitats where the host evolve and the identification of signatures of local adaptation on the candidate genes has been overlooked. To fill these gaps and dissect the genetic architecture driving adaptive plant-microbiota interactions, we adopted a Genome-Environmental-Association (GEA) analysis on 141 whole-genome sequenced natural populations of *Arabidopsis thaliana* characterized *in situ* for their leaf and root bacterial communities and a large range of environmental descriptors (i.e. climate, soil and plant communities). Across 194 microbiota traits, a much higher fraction of among-population variance was explained by the host genetics than by ecology, with the plant neighborhood as the main ecological driver of microbiota variation. Importantly, the relative importance of host genetics and ecology expressed a phylogenetic signal at the family and genus level. In addition, the polygenic architecture of adaptation to bacterial communities was highly flexible between plant compartments and seasons. Relatedly, signatures of local adaptation were stronger on QTLs of the root microbiota in spring. Finally, we provide evidence that plant immunity, in particular the *FLS2* gene, is a major source of adaptive genetic variation structuring bacterial assemblages in *A. thaliana*.

## Introduction

To cope with human population growth and current social requests, a more eco-efficient, sustainable and environmentally friendly agriculture is an urgent need (Keating et al. 2010). In the global change context, both crops and wild plant species face extreme and largely unpredictable abiotic stresses (such as heat waves) as well as an increase in the number and severity of epidemics (Bebber 2015; Desaint et al. 2021). Altogether, this calls for concrete interventions improving the potential of plants to cope with multiple abiotic and biotic stresses.

The plant microbiota is defined as a set of microorganisms of a particular host compartment (i.e. rhizosphere, roots, stem, leaves, flowers etc.). Often referred to as the second host genome in the context of the holobiont/hologenome theory (Rosenberg and Zilber-Rosenberg 2018), the plant microbiota mainly originates from the soil compartment, even if a non-negligible fraction of microbes also originates from the aerial sphere (Müller et al. 2016). Plant-associated microbes are crucial for plant health because they: i) mobilize and make accessible essential nutrients (e.g. nitrogen, phosphate etc.), ii) provide resistance to abiotic stresses (such as drought), and iii) confer direct (production of antimicrobial components) or indirect (elicitation of immune defense) pathogen protection (Berendsen et al. 2012; Bulgarelli et al. 2013; Pieterse et al. 2014; Jacoby et al. 2017; Escudero-Martinez and Bulgarelli 2019; Trivedi et al. 2020; Glick and Gamalero 2021; Bai et al. 2022). The plant microbiota is therefore a promising lever to develop innovative eco-friendly agro-ecosystems (Busby et al. 2017; Toju et al. 2018; Mitter et al. 2019).

While numerous studies reported the strong influence of abiotic (i.e. climate, physico-chemical agronomic properties) (Müller et al. 2016; Fitzpatrick et al. 2020) and biotic (i.e. presence of herbivores and neighboring plants) (Humphrey and Whiteman 2020; Meyer et al. 2022) factors on plant microbiota diversity and composition, there was a growing interest during the last decade to estimate the effect of plant genetics on microbiota (Bergelson, Brachi, et al. 2021). Two main approaches have been adopted to unravel the genetic and molecular plant mechanisms controlling microbiota assembly. The first and most common approach is based on the use of artificial mutations, including mutant and transgenic lines (Bergelson, Brachi, et al. 2021). By testing 218 artificial lines in 48 studies conducted on crops and wild species, several pathways affecting microbial assembly were identified and include (i) external and internal physical barriers in both the leaf (e.g. wax and cuticle) and root (e.g. suberin and lignin) compartments (Salas-González et al. 2021), (ii) Pathogen-Associated Molecular Pattern (PAMP)-triggered immunity (PTI) that prevents dysbiosis by keeping commensal microbes at a low absolute abundance (Chen et al. 2020), (iii) hormonal pathways related to salicylic acid, jasmonic acid, ethylene and strigolactones (Lebeis et al. 2015), (iv) mineral nutrient homeostasis (Zhang et al. 2019), which may requires a fine coordination with physical barriers (Salas-González et al. 2021) and immunity (Castrillo et al. 2017), (v) excretion of plant secondary metabolites in rhizosphere, roots or flowers (Huang et al. 2019), and (vi) symbiosis (Wang et al. 2020).

The second approach exploits natural genetic variation segregating among or within plant species (Bergelson, Brachi, et al. 2021). While the importance of host genetics in shaping natural variation on microbial communities has been a long-lasting debate (Roux and Bergelson 2016), an ever-increasing number of studies reported significant microbiota differences between closely related species or among genotypes within a given species (when grown in the same environment), with typical values for the magnitude of these differences ranging from 5% to 30% (Schlaeppi et al. 2014; Bergelson, Brachi, et al. 2021). Following the detection of significant heritability estimates, seven genome-wide association studies (GWAS) using microbial community descriptors as plant traits have been performed in *Arabidopsis thaliana* (Horton et al. 2014; Bergelson et al. 2019; Brachi et al. 2022), maize (Walters et al. 2018), rice (Roman-Reyna et al. 2020), sorghum (Deng et al. 2021) and switchgrass (VanWallendael et al. 2022). These GWAS revealed a highly polygenic architecture, suggesting a control of natural microbiota assembly by an extensive number of Quantitative Trait Loci (QTLs) with a small effect. A similar result was recently obtained with traditional linkage mapping performed in barley (Escudero-Martinez et al. 2022) and tomato (Oyserman et al. 2022).

While informative, the number of genetic association studies that report signatures of local adaptation on QTLs associated with microbial communities remains scarce, not to say absent. In addition, because the relative effect of host genetics on the microbiota can highly depend on the plant habitat, in particular the inoculum source (e.g. agricultural soil) (Robertson-Albertyn et al. 2017; Hubbard et al. 2018; Fabiańska et al. 2020), setting up genetic association studies in a common garden can bring to partial conclusions on the host genetics controlling for microbiota (Oyserman et al. 2020). One approach to tackle these issues is to conduct Genome-Environment Association (GEA) analysis. With the goal of identifying genetic variants associated with ecological variation across tens to hundreds of natural populations, GEA analysis is a powerful genome scan method to identify genes potentially involved in adaptive processes (De Mita et al. 2013). The development of next-generation sequencing (NGS) technologies combined with the availability of public databases on abiotic variables, in particular climatic variables, resulted in a recent burst of GEA studies reporting in plants the identification of adaptive QTLs to abiotic variation, from a worldwide scale (Hancock et al. 2011; Lasky et al. 2015; Bay et al. 2017; López-Hernández and Cortés 2019) to a regional scale (Pluess et al. 2016; Frachon et al. 2018). Although much less applied on biotic factors, a GEA analysis on plant community descriptors revealed a high degree of biotic specialization of *Arabidopsis thaliana* to members of its plant interaction network at the genetic level (Frachon et al. 2019). In addition, the genetic architecture of local adaptation to plant community diversity and composition was not predictable from the genetic architecture of local adaptation to the abundance of individual plant companion species (Frachon et al. 2019).

In this study, by combining microbial community ecology and population genomics, we adopted a GEA approach to establish a genomic map of adaptation to bacterial communities of the leaf and root compartment of 141 whole-genome sequenced natural populations of *A. thaliana* located south-west of France (Bartoli et al. 2018; Frachon et al. 2018). Because plant-associated microbes rapidly change within the host life cycle (Copeland et al. 2015; Beilsmith et al. 2021), bacterial communities were characterized in fall and spring, thereby allowing testing whether the strength of adaptation to bacterial communities differs between seasons (Bartoli et al. 2018). We also tested whether the strength of adaptation differs between microbiota and pathobiota (i.e. the ensemble of potential phytopathogens). To control for the confounding effects of the abiotic environment on microbiota and pathobiota, the 141 natural populations of *A. thaliana* were characterized for a set of 17 biologically meaningful climate and soil variables (Frachon et al. 2018; Frachon et al. 2019). Because companion species can strongly shape the microbial communities of a focal plant species (Geremia et al. 2016; Meyer et al. 2022), we also controlled for the confounding effects of 49 plant community descriptors (Frachon et al. 2019).

## Results & Discussion

A set of 168 natural populations of *A. thaliana* inhabiting contrasting ecological habitats in the south-west of France were whole-genome sequenced using a Pool-Seq approach, resulting in the identification of 4,781,661 SNPs (Frachon et al. 2018). In this study, we focused on 141 of these natural populations of *A. thaliana* that were characterized for leaf and root bacterial communities - using a *gyrB* based metabarcoding approach (Bartoli et al. 2018) - and a set of six climate variables, 14 soil physico-chemical variables and 49 descriptors of plant communities (supplementary table 1, Data Sets 1-8) (Frachon et al. 2018; Frachon et al. 2019). Importantly, the 141 populations strongly differed in their main germination cohort in autumn 2014 (early November vs. early December) (Bartoli et al. 2018). We therefore defined three seasonal groups, hereafter named (i) “fall” corresponding to 73 populations collected in November/December 2014, (ii) “spring (November)” corresponding to 72 populations already sampled in fall and additionally sampled in early-spring (February/March 2015), and (iii) “spring (December)” corresponding to 66 populations only sampled in early-spring (February/March 2015) (table 1, supplementary table 1).

**Table 1.** Number of bacterial community descriptors investigated with GEA analysis for each of the six ‘plant compartment × seasonal group’ combination. Numbers in brackets for the leaf and root compartment correspond to the number of populations. For community composition, only PCoA axes exhibiting significant variation among populations were considered in this study.

| Category | Community descriptors | fall |  | spring (November) |  | spring (December) |  |
| --- | --- | --- | --- | --- | --- | --- | --- |
|  |  | leaf (n = 73) | root (n = 73) | leaf (n = 69) | root (n = 72) | leaf (n = 66) | root (n = 66) |
| microbiota | richness | 1 | 1 | 1 | 1 | 1 | 1 |
|  | Shannon index | 1 | 1 | 1 | 1 | 1 | 1 |
|  | composition (PCoA axis) | 2 | 2 | 2 | 2 | 2 | 2 |
|  | presence/absence OTUs | 33 | 15 | 31 | 21 | 33 | 13 |
| pathobiota | richness | 1 | 1 | 1 | 1 | 1 | 1 |
|  | Shannon index | 1 | 1 | 1 | 1 | 1 | 1 |
|  | composition (PCoA axis) | 2 | 1 | 2 | 1 | 2 | 1 |
|  | presence/absence OTUs | 1 | 0 | 1 | 0 | 1 | 0 |

For each ‘plant compartment × seasonal group’ combination, we focused for both microbiota and pathobiota (i.e. ensemble of pathogenic lineages), on descriptors of community diversity (richness and Shannon index) and community composition (approximated by the two first principal components from a simple unconstrained PCoA) (Bartoli et al. 2018), as well as the presence/absence of the most prevalent OTUs, resulting in a total of 194 descriptors of bacterial communities (table 1). To fine map QTLs associated with microbiota/pathobiota traits, we combined a Bayesian hierarchical model (BHM) explicitly accounting for the scaled covariance matrix of population allele frequencies (Ω), which makes the analyses robust to complex demographic histories (Gautier 2015), with a local score (LS) approach allowing the detection of QTLs with small effects (Bonhomme et al. 2019). The efficiency of the local score approach was demonstrated in GWAS conducted in *A. thaliana*, with the fine mapping down to the gene level and the functional validation of four QTLs associated with quantitative disease resistance to the bacterial pathogen *Ralstonia solanacearum* (Aoun et al. 2020; Demirjian et al. 2022). In addition, the local score approach was successfully applied in a recent GWAS on leaf bacterial communities characterized on 200 Swedish accessions of *A. thaliana* grown in four native habitats in Sweden (Brachi et al. 2022).

### GEA revealed a polygenetic architecture of adaptation to bacterial communities

Our Bayesian hierarchical model – local score (BHM-LS) combined approach successfully detected QTLs associated with diversity and composition of bacterial communities and the presence/absence of a particular OTU. For instance, a neat association peak was detected at the end of chromosome 5 and the beginning of chromosome 1 for variation in microbiota diversity and composition in the leaf compartment of the ‘spring (November)’ seasonal group, respectively (fig. 1A). Similarly, a neat peak of association was detected at the beginning of chromosome 3 for the presence/absence of *Pseudomonas viridiflava*, one of the most prevalent and abundant bacterial pathogens identified in natural populations of *A. thaliana* in several geographical regions (Karasov et al. 2014; Bartoli et al. 2018; Karasov et al. 2018) (fig. 1A).

**Figure 1.**
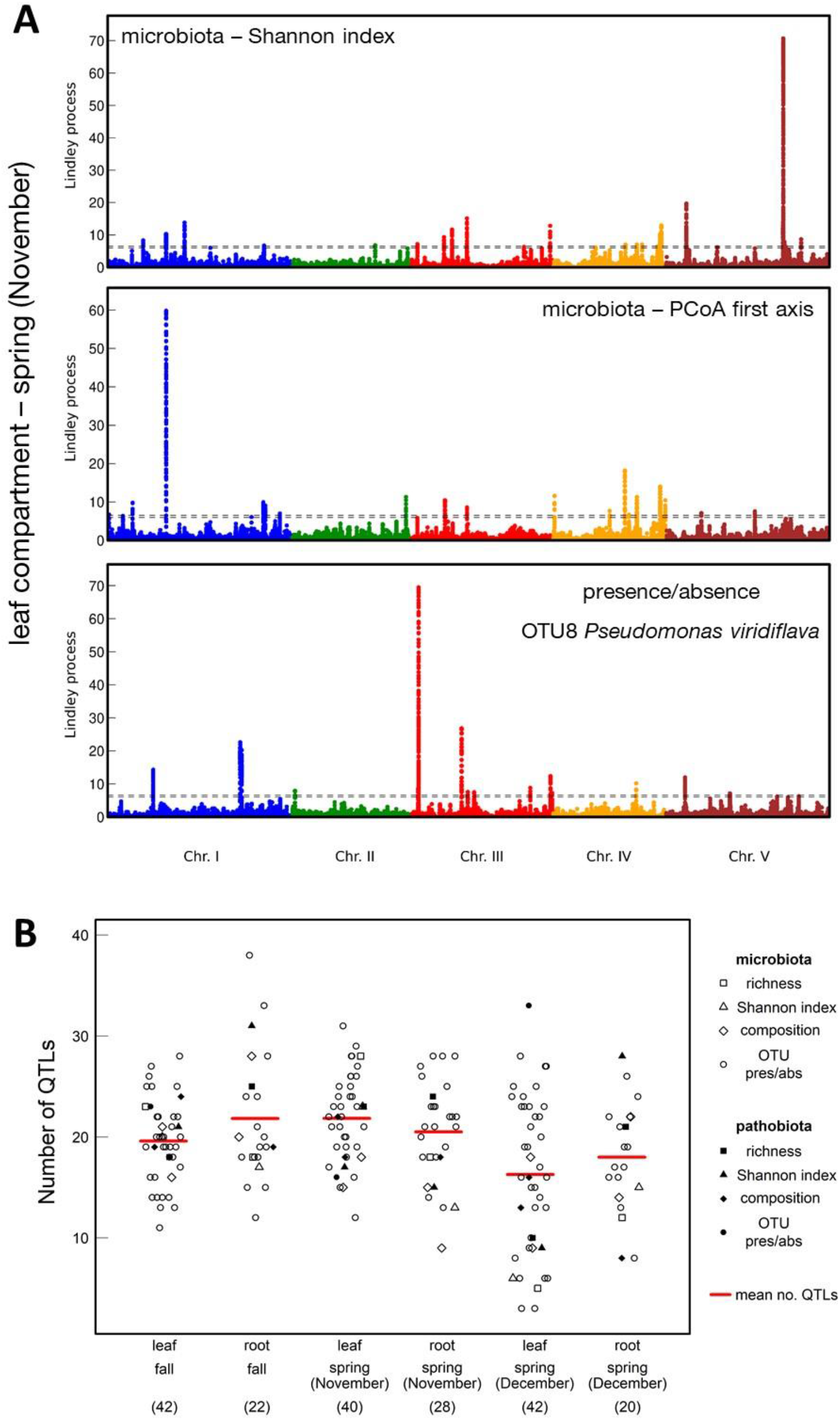
The genetic architecture of microbiota/pathobiota. (A) Manhattan plots illustrating the power of a BMH-LS to describe the genetic architecture of microbiota/pathobiota diversity, microbiota/pathobiota composition and the presence/absence of a particular OTU. The *x*-axis indicates along the five chromosomes, the physical position of the 1,470,777 SNPs considered for the leaf compartment in the ‘spring (November)’ seasonal group. The *y*-axis corresponds to the values of the Lindley process (local score method with a tuning parameter ξ = 2). The dashed lines indicate the minimum and maximum of the five chromosome-wide significance thresholds. (B) Jitter plots illustrating the diversity in the number of QTLs among microbiota/pathobiota traits within each ‘plant compartment × seasonal group’ combination.

Overall, our study revealed a highly polygenic architecture for most of the 194 descriptors of bacterial communities, with the detection of on average ∼19.6 QTLs per descriptor (median = 20, min = 3, max = 38) (fig. 1B). This is in line with the polygenic architecture reported in GWAS conducted on microbial communities (Horton et al. 2014; Walters et al. 2018; Bergelson et al. 2019; Roman-Reyna et al. 2020; Deng et al. 2021; Brachi et al. 2022; VanWallendael et al. 2022). Differences in the number of QTLs among the six ‘plant compartment × seasons’ were barely significant (Generalized Linear Model, GLM, *F* = 2.30, *P* = 0.0471). No differences in the number of QTLs was found between microbiota and pathobiota traits (GLM, *F* = 0.50, *P* = 0.8063) (fig. 1B).

### A non-negligible fraction of QTLs of microbiota/pathobiota were associated with variation in abiotic environment and plant communities

To disentangle GEA signals for a given bacterial community trait from GEA signals for abiotic variables and descriptors of plant communities, we applied BHM-LS on 69 climate variables, soil physico-chemical properties and plant community descriptors (supplementary table 1, Data Set 2-7). A non-negligible fraction of top SNPs associated with microbiota/pathobiota traits were common to variation in abiotic environment and plant communities, with on average ∼15.7% of top SNPs associated with microbiota/pathobiota traits being common with ecological variables (median = 13.2%, min = 0%, max = 84%) (fig. 2A). We may however caution that these values certainly represent lower-bound estimates due to the undefined number of ecological variables acting on natural populations of *A. thaliana*.

**Figure 2.**
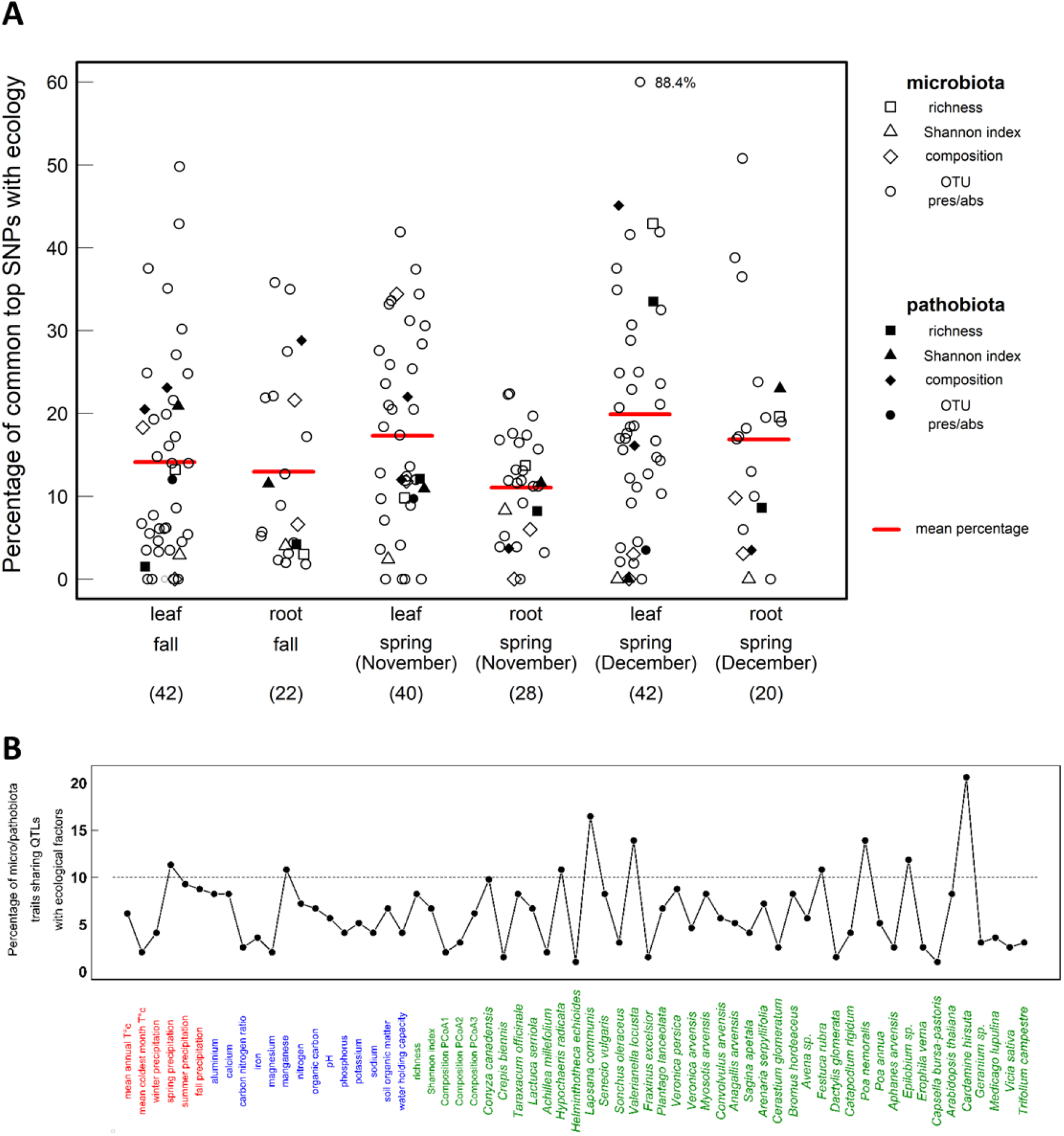
Microbiota/pathobiota-ecology interactions at the genetic level. (A) Percentage of top SNPs associated with microbiota/pathobiota traits that are common to top SNPs associated with ecological variables (including climate variables, soil physico-chemical properties and descriptors of plant communities) for each ‘plant compartment × seasonal group’ combination. (B) Percentage of microbiota/pathobiota traits (n = 194) sharing QTLs with each ecological factor among the six ‘plant compartment × seasonal group’ combinations. Climate variables, soil physico-chemical properties and descriptors of plant communities are represented in red, blue and green, respectively. The dashed black line corresponds to the threshold of 19 microbiota/pathobiota traits, i.e. ∼10% of the total number of microbiota/pathobiota traits.

Seven out of the nine ecological variables sharing top SNPs with more than 10% of microbiota/pathobiota traits correspond to the presence/absence of companion plant species (fig. 2B), which is in line with the neighborhood effects (also known as associational effects) on microbial transmission (Worrich et al. 2019; Meyer et al. 2022), particularly well-documented for bacterial pathogens (Parker et al. 2015; Makiola et al. 2022). For instance, a neat association peak located on chromosome 3 was common between the presence/absence of OTU6 - the Plant-Growth Promoting Bacteria (PGPB) *Pseudomonas siliginis* (Ramirez-Sanchez et al. 2022a) - in the root compartment in fall and the presence/absence of *Cardamine hirsuta*, a closely related to *A. thaliana* (Hay and Tsiantis 2016) (fig. 3A). A neat association peak located on chromosome 1 was common between the compositions in bacterial pathogens in the leaf compartment in fall and the richness in companion plant species (fig. 3B).

**Figure 3.**
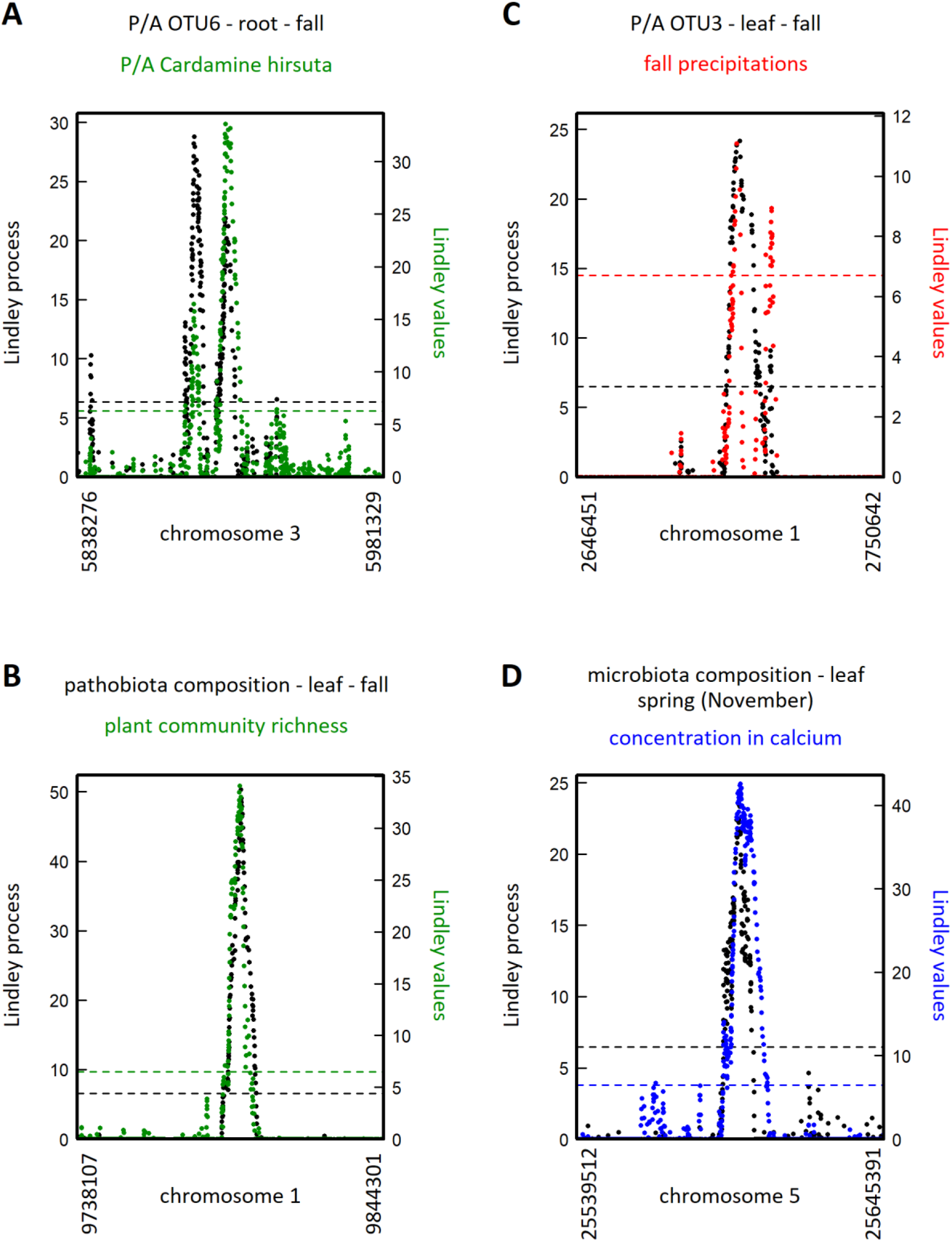
Zoom spanning four QTL regions common between microbiota/pathobiota traits and ecological variables. (A) Overlapping QTLs between the presence/absence of OTU6 in the root compartment in fall and the presence/absence of the plant species *Cardamine hirsuta*. (B) Overlapping QTLs between pathobiota composition in the leaf compartment in fall and plant community richness. Black and colored dots correspond to Lindley values for microbiota/pathobiota traits and ecological variables, respectively. The black and colored dashed lines indicate the corresponding chromosome-wide significance threshold for microbiota/pathobiota traits and ecological variables, respectively. (C) Overlapping QTLs between the presence/absence of OTU3 in the leaf compartment in fall and fall precipitations (mean value over the 2003-2013 period). (D) Overlapping QTLs between microbiota composition in the leaf compartment for the ‘spring (November)’ seasonal group and soil calcium concentrations.

In agreement with the impact of precipitation and drought in the phyllosphere microbiota (Zhu et al. 2022), the main climate variables sharing top SNPs with microbiota/pathobiota traits corresponds to precipitations (fig. 2A). For instance, a neat association peak located on chromosome 1 was common between the presence/absence of OTU3 in the leaf compartment in fall and precipitation in spring (fig. 3C). On the other hand, manganese and calcium concentrations are the main soil physico-chemical properties sharing top SNPs with microbiota/pathobiota traits (fig. 2A). For instance, microbiota composition in the leaf compartment in the ‘spring (November)’ seasonal group and soil calcium concentration shared a neat association peak located on chromosome 5 (fig. 3D). Soil calcium concentration was found to impact bacterial community structures both in soils (Neal and Glendining 2019; Tang et al. 2019) and in plants (Li et al. 2018; Mittelstrass et al. 2021), whereas soil manganese concentration was already suggested as a main driver of root microbial communities in *A. thaliana* at the continental scale (Thiergart et al. 2020).

For each seasonal group, the percentage of common SNPs between microbiota/pathobiota and ecology was on average higher in the leaf compartment than in the root compartment (fig. 2A), albeit not significant (GLM, *F* = 2.06, *P* = 0.1526). No differences was detected between microbiota and pathobiota descriptors (GLM, *F* = 0.06, *P* = 0.8082) (fig. 2A).

### The relative importance of host genetics and ecology in explaining microbiota/pathobiota variation expressed a phylogenetic signal across bacterial OTUs

In order to teasing apart the relative importance of host genetics and ecology in explaining microbiota/pathobiota variation, we run a multiple linear regression model on each microbiota/pathobiota trait by considering (i) all the QTLs that were specific to the microbiota/pathobiota trait under consideration (i.e. not common with ecological variables) and (ii) each ecological variable sharing QTLs with the microbiota/pathobiota trait under consideration (see Material & Methods), and (iii) a proxy considering the potential effects of demographic history of *A. thaliana* south-west of France, in explaining microbiota/pathobiota variation.

Across the 194 microbiota/pathobiota traits, a much higher fraction of among-population variance was explained by host genetics (mean = 35.4%, median = 35.0%) than by ecology (mean = 5.2%, median = 3.8%) (fig 4A, Data Set 9). In line with the polygenic architecture, the total fraction of variance explained by host genetics resulted from multiple QTLs, each explaining on average 1.99% (supplementary fig. S1, Data Set 10). The small QTL effects detected by GEA are similar to the QTL effect sizes identified in GWAS and traditional linkage mapping studies on microbiota (Bergelson, Brachi, et al. 2021; Oyserman et al. 2022), albeit a larger range of QTL effects was identified in our study. For instance, the top SNP located on position 19,335,417bp on chromosome 2 (SNP 2_19335471), SNP 1_12691514 and SNP 5_5191015 explained a substantial fraction of the presence/absence of OTU4 (root compartment in fall), microbiota diversity (leaf compartment in spring (December)) and pathobiota composition (leaf compartment in fall), respectively (fig. 4B, 4C and 4D, Data Set 10).

**Figure 4.**
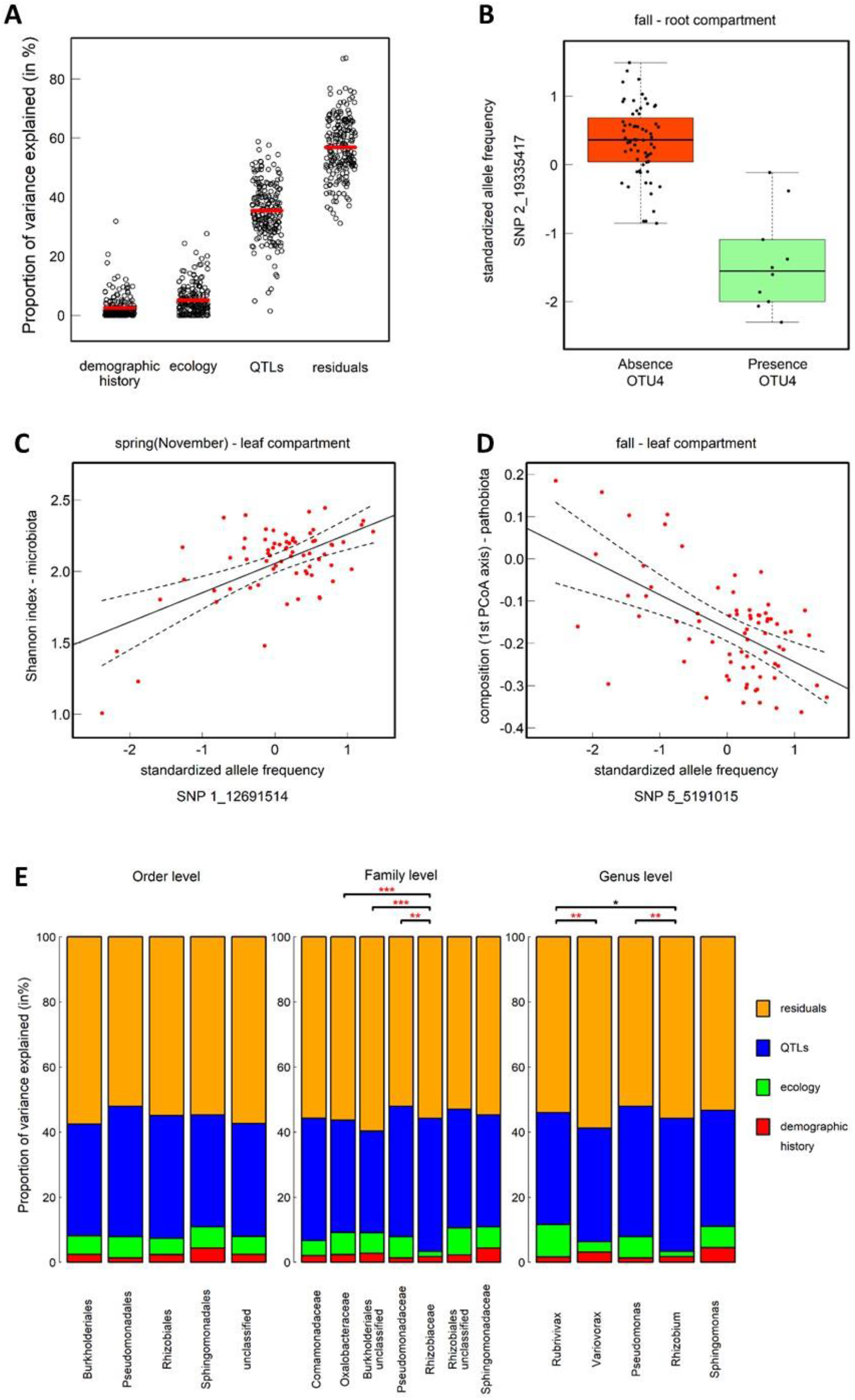
Relative importance of host genetics and environment in explaining variation of microbiota/pathobiota traits. (A) Cumulative percentage of variance explained (PVE) of microbiota/pathobiota traits by the demographic history of *A. thaliana*, host genetics with QTLs specific to microbiota/pathobiota traits and ecology (including climate variables, soil physico-chemical properties and descriptors of plant communities). Residuals correspond to the percentage of variance unexplained by the three other categories of variables. For each category, one dot correspond to one of the 194 microbiota/pathobiota traits. (B) Box-plot illustrating the relationship between the standardized allele frequencies (corrected for the effect of population structure) of a SNP located at position 19,335,417 on chromosome 2 and the presence of OTU4 in the root compartment in fall. (C) Relationship between the standardized allele frequencies (corrected for the effect of population structure) of a SNP located at position 12,691,514 on chromosome 1 and Shannon index of the microbiota in the leaf compartment of the ‘spring (November)’ seasonal group. (D) Relationship between the standardized allele frequencies of a SNP located at position 5,191,015 on chromosome 5 and pathobiota composition in the leaf compartment in fall. (E) Relative importance of PVE by demographic history, host genetics, ecology and residuals, among OTUs at the order, family and genus level. * *P* < 0.05, ** *P* < 0.001, *** *P* < 0.001. Red asterisks indicate significant *p*-values after correcting for multiple testing with a false discovery rate (FDR) at a nominal level of 5%.

Amongst ecological variables, the variance in microbiota/pathobiota traits was on average more explained by plant community descriptors (mean = 3.75%) than by climate variables (mean = 0.66%) and soil physico-chemical properties (mean = 0.78%), highlighting the plant neighborhood as the main ecological driver of microbiota/pathobiota of *A. thaliana* populations located south-west of France. A very small fraction of microbiota/pathobiota variation was on average explained by the demographic history of *A. thaliana* (mean = 2.52%, median = 1.23%) (supplementary fig. 1, Data Set 9). Across the 194 microbiota/pathobiota traits, a substantial fraction of variance remained unexplained (mean = 56.9%, median = 56.4%) (supplementary fig. S1, Data Set 9). This unexplained variance may originate from (i) QTLs with very small effect that remains undetected due to a lack of power given the number of populations used for GEA analysis (table 1), (ii) uncharacterized explanatory ecological variables, as previously mentioned, and/or (iii) stochastic processes of dispersal and drift that can drastically alter community structure (Wagner 2021).

The relative importance of host genetics, ecology and demographic history in explaining microbiota/pathobiota variation was similar between leaf and root compartments, between microbiota and pathobiota, and between the three seasonal groups (supplementary fig. 2, supplementary table 2). Importantly, the relative importance of host genetics, ecology and demographic history expressed a phylogenetic signal at the family and genus level (fig. 4E). For instance, the relative importance of host genetics (in comparison with ecology) in explaining variance in microbiota/pathobiota OTUs was significantly higher for OTUs of the *Rhizobium* genus than for OTUs of the *Pseudomonas* genus (fig. 4E). A phylogenetic signal at the order level was previously observed in GWAS conducted on the rhizospheric microbiota of sorghum and the leaf microbiota of maize, with heritable microbes that phylogenetically clustered (Walters et al. 2018; Deng et al. 2021). Altogether, these results suggest that some taxonomic groups necessitate more genetic discrimination among genotypes of *A. thaliana* than others, thereby providing candidate bacterial taxa to be investigated for genomic signals of co-evolution with *A. thaliana* through, for example, free-phenotyping co-GWAS (Bartoli and Roux 2017).

### The genetic architecture of adaptation to bacterial communities is highly flexible between plant compartments and seasons

After retrieving candidate genes located in the vicinity of the top SNPs specific to the microbiota/pathobiota trait under consideration (i.e. not common with ecological variables), we observed a highly flexible genetic architecture between the six ‘plant compartment × seasonal group’ combinations, with 76.1% of unique candidate genes being specific to a ‘plant compartment × seasonal group’ combination (fig. 5A, Data Set 11). Most of the remaining candidate genes were either specific to the leaf compartment and common between the three seasonal groups, or specific to a given seasonal group and common between the leaf and root compartments (fig. 5A). On the other hand, very few candidate genes were specific to the root compartment and common between the three seasonal groups (fig. 5A).

**Figure 5.**
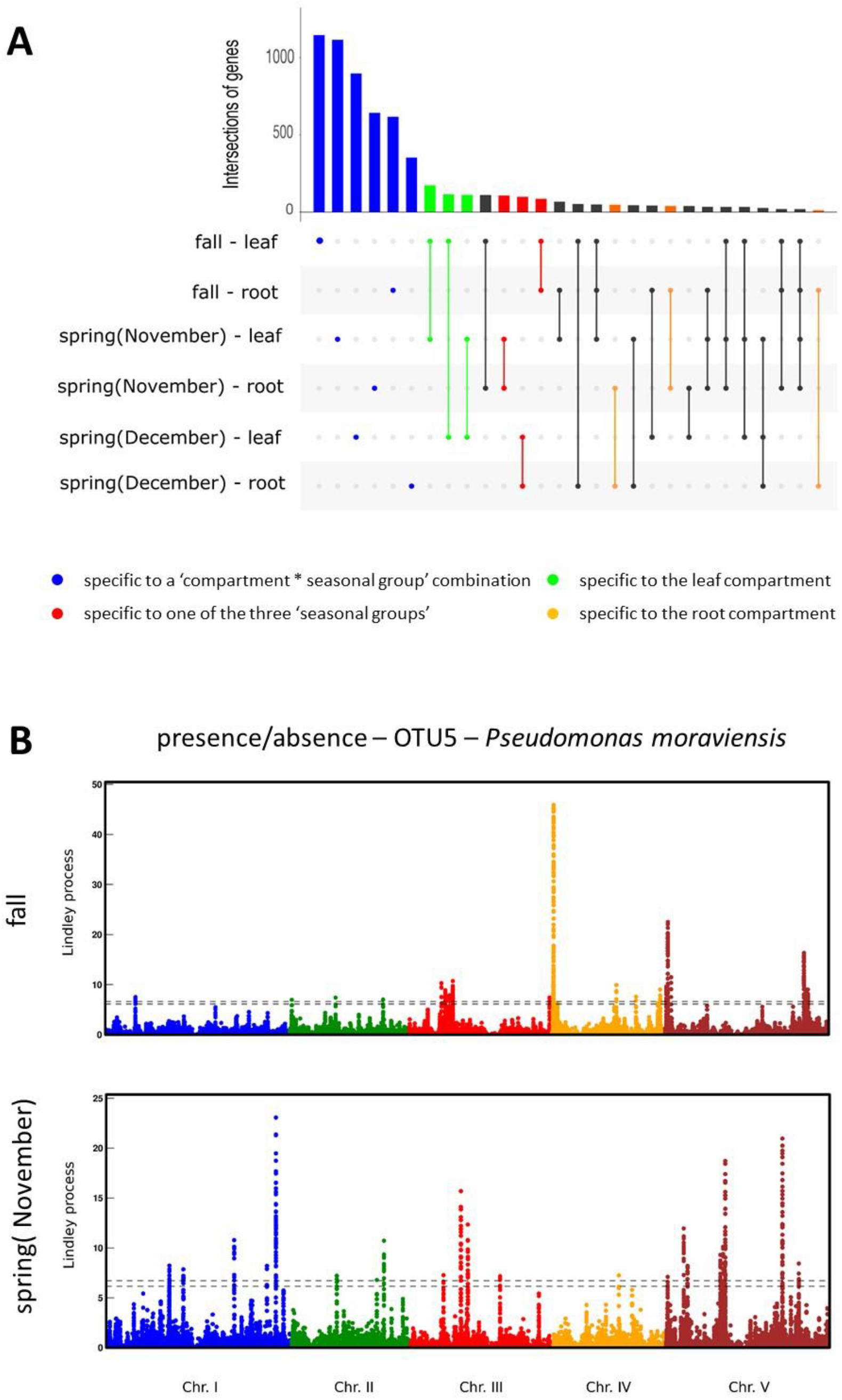
Flexibility of the genetic architecture between the six ‘plant compartment × seasonal group’ combinations. (A) An UpSet plot showing the intersections of the lists of unique candidate genes associated with microbiota/pathobiota variation, each list corresponding to each ‘plant compartment × seasonal group’ combination. The number of candidate genes that are specific to a single ‘plant compartment × seasonal group’ combination and common between at least two ‘plant compartment × seasonal group’ combinations, are represented by single blue dots and dots connected by a solid line, respectively. ‘fall – leaf’: number of unique candidate genes *N* = 1918, ‘fall – root’: *N* = 1052, ‘spring (November) – leaf’: *N* = 1885, ‘spring (November) – root’: *N* = 1156, ‘spring (December) – leaf’: *N* = 1457, ‘spring (December) – root’: *N* = 705. (B) Manhattan plot illustrating the flexibility of genetic architecture associated with the presence/absence of the OTU5 in the root compartment between fall and spring. The x-axis indicates along the five chromosomes, the physical position of the 1,392,959 SNPs and 1,382,414 SNPs considered for the root compartment in the ‘fall’ and ‘spring (November)’ seasonal groups, respectively. The y-axis correspond to the values of the Lindley process (local score method with a tuning parameter ξ = 2). The dashed lines indicate the minimum and maximum of the five chromosome-wide significance thresholds.

All GWAS performed previously on plant microbial communities were conducted on one specific plant compartment (Walters et al. 2018; Roman-Reyna et al. 2020; Deng et al. 2021; Brachi et al. 2022; VanWallendael et al. 2022), with the exception of one GWAS conducted on worldwide accessions of *A. thaliana* at the leaf (Horton et al. 2014) and root (Bergelson et al. 2019) levels. Similar to our results, this latter GWAS showed a small overlap between the leaf and root compartments in candidate genes associated with descriptors of bacterial and fungal communities (Bergelson et al. 2019), which is in line with the adaptive differences in microbial community diversity and composition observed among plant niches, from rhizosphere soils to plant canopies (Müller et al. 2016; Cregger et al. 2018).

As illustrated with OTU5 corresponding to the PGPB *Pseudomonas moraviensis* (Ramirez-Sanchez et al. 2022a), we observed a strong flexibility of genetic architecture between seasons for a specific microbiota/pathobiota trait (fig. 5B). Such an observation is reminiscent of the strong genetic variation in seasonal microbial community succession observed in diverse plant species including *A. thaliana* (Copeland et al. 2015; Bartoli et al. 2018; Beilsmith et al. 2021; VanWallendael et al. 2022) and the dynamics of genetic architecture along the infection stages when *A. thaliana* accessions were challenged with the bacterial pathogen *Ralstonia solanacearum* (Aoun et al. 2017; Aoun et al. 2020; Demirjian et al. 2022).

The identity of candidate genes strongly differs between the two seasonal groups ‘spring (November)’ and ‘spring (December)’ in both plant compartments, thereby suggesting an effect of germination timing on the interplay between host genetics and microbiota/pathobiota. While germination timing was found to influence natural selection on life-history traits in *A. thaliana* (Donohue 2002; Donohue et al. 2005), the effect of germinating timing on microbiota/pathobiota assemblages has been seldom reported and deserves further investigations.

Beyond a highly flexible genetic architecture between plant compartments and seasons, we also observed a highly flexible genetic architecture among microbiota/pathobiota traits for each ‘plant compartment × seasonal group’ combination (supplementary fig. 3). For instance, for the pathobiota in the leaf compartment in the ‘spring (December)’ seasonal group, the identity of candidate genes strongly differs between community diversity, community composition and the presence/absence of *P. syringae* (supplementary fig. 4). As previously observed in GWAS and GEAS conducted on plant-plant interactions in *A. thaliana* (Baron et al. 2015; Frachon et al. 2019; Libourel et al. 2021), these results suggest a high degree of biotic specialization of *A. thaliana* to members of its bacterial interaction network, as well as the genetic ability of *A. thaliana* to interact simultaneously with multiple bacterial members.

Altogether, in line with the ever-changing complexity of biotic interactions observed in nature (Bergelson, Kreitman, et al. 2021), our study reinforces the need to conduct association genetic studies on diverse plant compartments and seasons to obtain a full picture of the host genetics controlling natural variation of microbiota/pathobiota.

### The strength of signatures of local adaptation on QTLs of microbiota/pathobiota depends on plant compartment and season

By definition, GEA allows identifying genetic loci under local adaptation. However, in order to support that the loci identified by our GEA analysis have been shaped by natural selection, we additionally tested whether the top SNPs specific to microbiota/pathobiota traits were enriched in a set of SNPs subjected to adaptive spatial differentiation. To do so, we first performed for each ‘plant compartment × seasonal group’ combination a genome-wide selection scan by estimating a Bayesian measure of genetic differentiation (XtX) among the natural populations of *A. thaliana*. For a given SNP, XtX measures the variance of the standardized population allele frequencies, which is corrected for the genome-wide effects of confounding demographic evolutionary forces (Gautier 2015). The 0.5% upper tail of the spatial differentiation distribution displayed a significant enrichment (up to 34.2) for top SNPs of almost two-thirds of the microbiota/pathobiota traits (fig. 6A, Data Set 12). For instance, a strong signature of local adaptation was identified on SNPs located in the 5’ region of *MYB15* (fig. 6B), a transcription factor involved in the coordination of microbe-hots homeostasis in *A. thaliana* (Ma et al. 2021).

**Figure 6.**
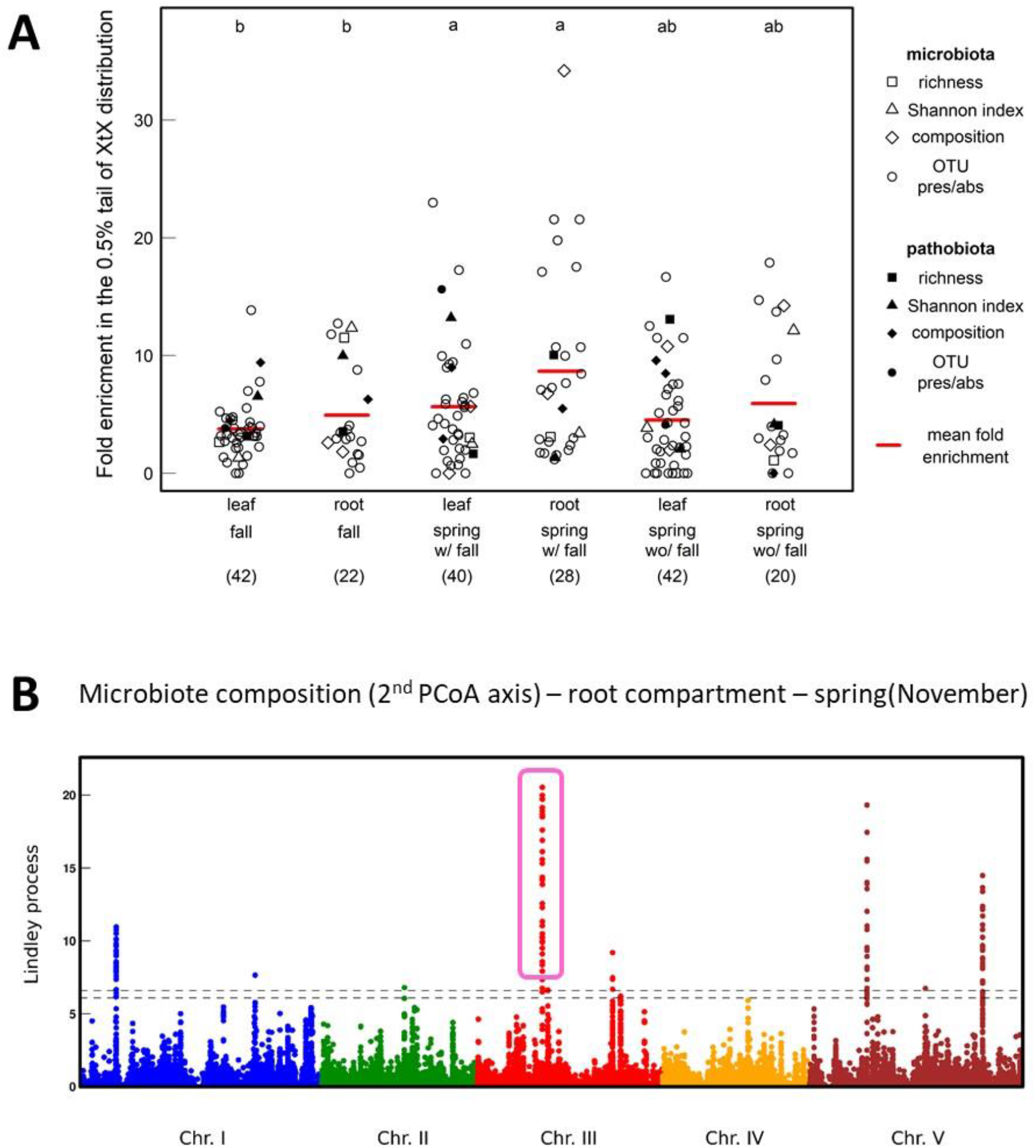
Signatures of local adaptation acting on the genetics of microbiota/pathobiota variation. (A) Fold enrichment of the top SNPs in the 0.5% tail of the genome-wide spatial differentiation (XtX) distribution. Each dot corresponds to one of the 194 microbiota/pathobiota descriptors. Different upper letters indicate different groups according to the ‘plant compartment × seasonal group’ combination after a Tukey correction for multiple pairwise comparisons. (B) Manhattan plot highlighting a QTL associated with root microbiota composition, located in the middle of chromosome 3 and presenting a strong signature of local adaptation. The *x*-axis indicates along the five chromosomes, the physical position of the 1,382,414 SNPs considered for the root compartment in the ‘spring (November)’ seasonal group. The y-axis correspond to the values of the Lindley process (local score method with a tuning parameter ξ = 2). The dashed lines indicate the minimum and maximum of the five chromosome-wide significance thresholds.

No differences in the mean fold enrichment for signatures of local adaptation was detected between microbiota and pathobiota traits (GLM, *F* = 0.44, *P* = 0.7260). However, the mean fold enrichment for signatures of local adaptation largely differed between the three seasonal groups (GLM, *F* = 5.06, *P* = 0.0072), with top SNPs presenting more signatures of local adaptation in spring than in fall when considering the same set of populations, i.e. populations from the ‘fall’ and ‘spring (November)’ seasonal groups (fig. 6A). Top SNPs of the ‘spring (December)’ seasonal group presented signatures of local adaptation that were intermediate between the two other seasonal groups, suggesting that germinating timing not only affects the genetic architecture underlying microbiota/pathobiota variation but also the strength of local adaptation acting on candidate genes.

In addition, the mean fold enrichment for signatures of local adaptation was significantly higher for the root compartment than for the leaf compartment (GLM, *F* = 6.11, *P* = 0.0143). Combined with the observation that the percentage of common SNPs between microbiota/pathobiota and ecology was on average higher in the leaf compartment than in the root compartment, the difference between the leaf and root compartments in the strength of local adaptation acting on candidate genes suggests a higher adaptive genetic control of the root microbiota than of the leaf microbiota.

### Cross validation of our GEA approach through results obtained from previous GWAS

A GEA approach suffers from the quasi-impossibility of measuring the entire set of ecological variables acting on natural populations of *A. thaliana*, thereby precluding the identification of all SNPs common between microbiota/pathobiota traits and ecological variables. To circumvent this issue, we therefore tested whether our list of candidate genes associated with microbiota/pathobiota traits significantly overlap with candidate genes identified in two GWAS conducted on *A. thaliana* in common gardens under ecologically relevant conditions. The first GWAS was conducted on 200 Swedish accessions characterized for the bacterial and fungal communities in the leaf compartment in the native habitat of four populations located in Sweden (Brachi et al. 2022). The list of 880 candidate genes located in the vicinity of the top SNPs associated with bacterial hubs in Sweden significantly overlapped with the lists of candidate genes identified by GEA for the leaf compartment, whatever the seasonal group (supplementary fig. 5A, Data Set 13). The fraction of candidate genes detected in Sweden and overlapping with the lists of candidate genes identified by GEA for the root compartment, was smaller and less significant for each seasonal group (supplementary fig. 5A).

The second GWAS was conducted on 162 whole-genome sequenced accessions of *A. thaliana* originating from 54 out of the 141 natural populations used in our study. Those accessions were challenged in field conditions with 13 bacterial strains belonging to seven of the 12 most abundant and prevalent leaf OTUs across our natural populations in south-west of France (Ramirez-Sanchez et al., 2022b). The resulting list of candidate genes showed a significant enrichment in diverse biological pathways, including cell, cell wall, development, hormone metabolism, secondary metabolism, signalling and transport, which is in line with the main enriched biological pathways identified in our study (supplementary fig. 5B, Data Set 14) and most of the main biological pathways (with the exception of symbiosis) identified by the use of artificial mutations and by previous GWAS conducted on *A. thaliana* (Bergelson, Brachi, et al. 2021).

Altogether, the similar lists of candidate genes and biological pathways detected between diverse mapping populations and using different approaches in association genetics, indicate a cross-validation of our GEA results by GWAS conducted in ecologically relevant conditions.

### *FLS2* as one of the main candidate genes controlling structure of bacterial communities

Merging the lists of candidate genes identified in our study and in two GWAs conducted in ecologically relevant conditions in the native area of *A. thaliana* (Brachi et al. 2022) led to the establishment of a short list of 50 core genes associated with natural variation of microbiota traits (Data Set 15). Of particular interest is the gene *FLAGELLIN-SENSITIVE 2* (*FLS2*) overlapping with 39 top SNPs in our study (Data Set 11). FLS2 encodes for a leucine-rich repeat receptor-like kinase first identified in the perception of the bacterial elicitor flagellin (Gómez-Gómez and Boller 2000) and then confirmed as key in microbe-associated molecular patterns (MAMP)-triggered immunity (MTI) (Stringlis and Pieterse 2021). Although *FLS2* detects flagellin from both pathogenic and beneficial bacteria by the 22-amino-acid N-terminal epitope flg22 (Stringlis et al. 2018), *FLS2* was never proposed as a candidate gene in GWAS of response to bacterial pathogens (https://aragwas.1001genomes.org/#/gene/AT5G46330), despite the identification of more than 100 amino acid changes in FLS2 among >1,000 worldwide accessions (Bai et al. 2022). Genetic variation at *FLS2* might have evolved to detect the substantial diversity of flg22 encoded by commensal bacteria (Colaianni et al. 2021; Bai et al. 2022). Accordingly, *FLS2* shaped the microbiota composition of *A. thaliana* rhizosphere when plant were grown on an agricultural soil (Fonseca et al. 2022). Assessing the allelic diversity of both *FLS2* and flg22 in our collection of 168 natural populations of *A. thaliana* (Frachon et al. 2018) and in our collection of >7,000 bacterial strains (Ramirez-Sanchez et al. 2022a), respectively, might reveal signatures of co-adaptation.

## Conclusion

The GEA study conducted here on microbiota/pathobiota traits and on an unprecedented number of ecological variables, is complementary to GWAS carried out previously on *A. thaliana*. In particular, we investigated (i) the level of flexibility of the genetic architecture between seasons and between *in situ* germination timings, (ii) the relative importance of host genetics and ecology in explaining microbiota/pathobiota variation, (iii) the identity of the ecological variables acting as putative selective agents on microbiota/pathobiota traits, and (iv) the variation between plant compartments and seasonal groups, in the strength of local adaptation acting on candidate genes. Importantly, no differences were observed between microbiota and pathobiota traits, thereby suggesting a similar adaptive genetic control of *A. thaliana* towards its pathogenic and non-pathogenic members, including members with already described beneficial effects (Ramirez-Sanchez et al. 2022).

The next avenue is then to apply our approach on fungal communities that have not been characterized on our set of natural populations yet. Establishing a genomic map of local adaptation to fungal communities might allow testing whether the genetic bases largely differ between bacterial and fungal communities, as previously documented in the two GWAS on *A. thaliana* (Horton et al. 2014; Bergelson et al. 2019; Brachi et al. 2022). It might also help to identify the adaptive genetic bases of common interactions among OTUs, including mutualism, antagonism, aggression and altruism, within and across kingdoms (He et al. 2021).

## Materials and Methods

### Descriptors of bacterial communities in leaves and roots

The bacterial communities of 1,903 leaf and root samples collected in fall and spring 2015 across 163 out of 168 natural populations of *A. thaliana* located south-west of France (Frachon et al. 2018), were characterized with a *gyrB*-based metabarcoding approach, leading to the identification of 278,833 OTUs (Bartoli et al. 2018). The deeper taxonomic resolution conferred by the *gyrB* gene allows distinguishing OTUs belonging to the pathobiota from OTUs belonging to the microbiota (Bartoli et al. 2018).

In this study, we considered 141 natural populations of *A. thaliana* for which a set of six climate variables, 14 soil physico-chemical variables and 49 descriptors of plant communities were available (see below), thereby resulting in 73, 73, 69, 72, 66 and 66 populations for the ‘fall – leaf’, ‘fall – root’, ‘spring (November) – leaf’, ‘spring (November) – root’, ‘spring (December) – leaf’ and ‘spring (December) – root’ combinations, respectively (table 1).

For each sample of *A. thaliana* collected in the 141 populations, we estimated the relative abundance (N° of reads per OTU / N° of total reads) of each of the 278,333 OTUs. For each ‘plant compartment × seasonal group’ combination, we then averaged for each OTU the corresponding relative abundances per population. For each OTU, populations with a mean relative abundance above and below (or equal to) 0.5% were scored as 1 (presence of the OTU) and 0 (absence of the OTU), respectively. In this study, for the purpose of statistical power in GEA analysis, we only kept OTUs present in more than seven populations, resulting in 34 OTUs, 15 OTUs, 32 OTUs, 21 OTUs, 34 OTUs and 13 OTUs investigated for GEA analysis for the ‘fall – leaf’, ‘fall – root’, ‘spring (November) – leaf’, ‘spring (November) – root’, ‘spring (December) – leaf’ and ‘spring (December) – root’ combinations, respectively (table 1). Similarly, for each ‘plant compartment × seasonal group’ combination, we averaged for both the microbiota and the pathobiota, estimates of community richness and Shannon index previously obtained in (Bartoli et al. 2018), as well as estimates of community composition using the coordinates of the samples on the two first axes of a principal coordinate analysis (PCoA) run on a Hellinger distance matrix based on the relative abundances of the 6,627 most abundant OTUs (Data Set 1) (Bartoli et al. 2018).

### Abiotic descriptors and descriptors of plant communities

A set of six climate variables, 14 soil physico-chemical properties and 49 descriptors of plant communities were available for 141 out of the 168 natural populations of *A. thaliana* located south-west of France (Data Sets 2-8). The climate variables corresponds to two variables related to temperature and four variables related to precipitation (Frachon et al. 2018). The 14 soil physico-chemical variables describe the main soil agronomic properties related to plant growth (Brachi et al. 2013; Frachon et al. 2019). The 49 descriptors of plant communities correspond to (i) estimates of richness and Shannon index (ii) estimates of community composition using the coordinates of the populations on the three first axes of a PCoA run on a Bray-Curtis dissimilarity matrix based on the relative abundance of the 44 most prevalent plant species and (iii) the presence/absence of the 44 most prevalent plant species, with the exception of *A. thaliana* for which estimates of the absolute abundance were kept (Frachon et al. 2019). In this study, for the purpose of statistical power in GEA analysis, we only kept companion plant species present in more than seven populations, resulting in 30, 29, 30, 31, 33 and 33 plant species investigated for GEA analysis for the ‘fall – leaf’, ‘fall – root’, ‘spring (November) – leaf’, ‘spring (November) – root’, ‘spring (December) – leaf’ and ‘spring (December) – root’ combinations, respectively (Data Sets 2-7).

### Genomic characterization and data filtering

As previously described in (Frachon et al. 2018), a representative picture of within-population genetic variation across the genome was previously obtained for the 168 populations located south-west of France, using a Pool-Seq approach based on the individual DNA extraction of ∼16 plants per population (min = 5 plants, max = 16 plants, mean = 15.32 plants, median = 16 plants). After bioinformatics analysis using the reference genome Col-0, the allele read count matrix (for both the reference and alternate alleles) was composed by 4,781,661 SNPs across the 168 populations (Frachon et al. 2018).

Following Frachon et al. (2018), for each ‘plant compartment × seasonal group’ combination, the matrix of population allele frequencies was trimmed according to five successive criteria : (i) removing SNPs with missing values in more than two populations, (ii) in order to take into account multiple gene copies in the populations that map to a unique gene copy in the reference genome Col-0, removing SNPs with a mean relative coverage depth across the populations above 1.5 after calculating for each population the relative coverage of each SNP as the ratio of its coverage to the median coverage (computed over all the SNPs), (iii) in order to take into account indels that correspond to either genomic regions present in Col-0 but absent in the populations or genomic regions present in the populations but absent in Col-0, removing SNPs with a mean relative coverage depth across the populations below 0.5, (iv) removing SNPs with a standard deviation of allele frequency across the populations below 0.004, and (v) in order to take into account bias in GEA analysis due to rare alleles (Bergelson and Roux 2010), removing SNPs with the alternative allele present in less than 10% of the populations. This SNP pruning resulted in a final number of 1,396,579 SNPs, 1,392,959 SNPs, 1,470,777 SNPs, 1,382,414 SNPs, 1,514,789 SNPs and 1,514,789 SNPs for the ‘fall – leaf’, ‘fall – root’, ‘spring (November) – leaf’, ‘spring (November) – root’, ‘spring (December) – leaf’ and ‘spring (December) – root’ combinations, respectively.

### Genome-Environment Association analysis

For each ‘plant compartment × seasonal group’ combination, a GEA analysis was performed between the set of pruned SNPs and each trait related to microbiota, climate, soil physico-chemical properties and plant communities, resulting in a total number of 530 traits, with 97 traits, 76 traits, 95 traits, 84 traits, 100 traits and 78 traits for the ‘fall – leaf’, ‘fall – root’, ‘spring (November) – leaf’, ‘spring (November) – root’, ‘spring (December) – leaf’ and ‘spring (December) – root’ combinations, respectively (supplementary table 1, Data Sets 2-7).

To identify significant ‘SNP-trait’ association s for each ‘plant compartment × seasonal group’ combination, we first run a Bayesian hierarchical model (i) explicitly accounting for the scaled covariance matrix of population allele frequencies (**Ω**), which makes the analyses robust to complex demographic histories (ii) dealing with Pool-Seq data and (iii) implemented in the program BayPass (Gautier 209715). Following (Frachon et al. 2019), the core model was used to evaluate the association between allele frequencies across the genome and the *n* traits. For each SNP, we estimated a Bayesian Factor (BF_is_ measured in deciban units) and the associated regression coefficient (Beta_is, *β*_i_) using an Importance Sampling algorithm (Gautier 2015). The full posterior distribution of the parameters was obtained based on a Metropolis–Hastings within Gibbs Markov chain Monte Carlo (MCMC) algorithm. A MCMC chain consisted of 15 pilot runs of 500 iterations each. Then, MCMC chains were run for 25,000 iterations after a 2500-iterations burn-in period. The *n* traits were scaled (*scalecov* option) so that μ = 0 and σ ^2^ = 1. Because of the use of an Importance Sampling algorithm, we repeated the analyses three times for each trait and averaged BF_is_ and *β*_i_ values across these three repeats. As previously performed in (Frachon et al. 2018), for each ‘plant compartment × seasonal group’ combination, we parallelized GEA analysis by dividing the full data set of pruned SNPs into 32 sub-data sets, each containing 3.125% of the total number of pruned SNPs taken every 32 SNPs across the genome.

As a second step, in order to better characterize the genetic architecture associated with ecological variation, the GEA results were reanalyzed by applying a local score approach (Bonhomme et al. 2019), which allows detecting significant genomic segments by accumulating the statistical signals from contiguous genetic markers such as SNPs (Fariello et al. 2017). In addition, this local score approach increases the power of detecting QTLs with small effect and narrows the size of QTL genomic regions (Fariello et al. 2017; Bonhomme et al. 2019). In a given QTL region, the association signal, through the *p*-values, will cumulate locally due to linkage disequilibrium between SNPs, which will then increase the local score (Bonhomme et al. 2019). Following (Libourel et al. 2021), in order to apply the local score approach on the GEA results, we first ranked each SNP based on the Bayes Factor values obtained across the genome (from the highest to the lowest values) for each trait. Then, each rank was divided by the total number of SNPs to obtain a *p*-value associated with each SNP. The local score approach was then implemented on these *p*-values to fine map genomic regions associated with traits. In this study, the tuning parameter ξ was fixed at 2 expressed in –log_10_ scale. Significant associations between SNPs and ecological variation were identified by estimating a chromosome-wide significance threshold for each chromosome (Bonhomme et al. 2019). The SNPs underlying the QTLs identified by the combined BMH-LS approach are hereafter named top SNPs.

### Estimating the relative importance of host genetics and environment in explaining variation of microbiota/pathobiota traits

To estimate the relative importance of (i) QTLs specific to microbiota/pathobiota traits, (ii) the demographic history of *A. thaliana*, and (ii) abiotic environment / plant communities in explaining variation of microbiota/pathobiota traits, we run the following multiple linear regression model under the *R* environment for each of the 194 microbiota/pathobiota traits:

Y_a,p,i…n,j…m,k_ = μ_a_ + population structure_p_ + QTL_i_ + … + QTL_n_ + ECOL_j_ + … + ECOL_m_ + ε_a,p,i…n,j…m,k_ Where Y is one of the 194 microbiota/pathobiota traits; *μ* is the overall mean; ‘population structure’ accounts for the effect of the demographic history of *A. thaliana* by using the coordinates of the populations on the first Principal Component axis (PC_genomic_ axis 1) resulting from the singular value decomposition of the scaled covariance matrix of population allele frequencies and explaining 96.4% of the genomic variation observed among the 168 populations located south-west of France (Frachon et al. 2018); ‘QTL’ accounts for the effect of host genetics by using the standardized allele frequencies (i.e. corrected for the effect of demographic history) (Gautier 2015) of the SNP with the highest BFis value for each QTL specific to the microbiota/pathobiota trait considered; ‘ECOL’ corresponds to values of ecological variables (climate variables, soil physico-chemical properties and descriptors of plant communities) for which QTLs were common to QTLs associated with microbiota/pathobiota traits; and ε is the residual term. For each microbiota/pathobiota trait, we obtained a percentage of variance explained (PVEs) by each model term (Data Set 10) and PVEs were then summed according to four categories, i.e. demographic history, host genetics specific to microbiota/pathobiota, ecology (including abiotic environment and plant communities) and residuals (Data Set 9).

To test whether the relative PVE by these four categories differs among OTUs at the order, family and genus taxonomic levels, we first averaged the PVE among OTUs belonging to a specific order, family or genus (only order, family or genus with at least four OTUs were considered). At each taxonomic level, we then applied a Chi-squared test on each pairwise comparison among OTUs. A Bonferroni correction was applied to control for multiple testing. A similar approach was applied to test whether the relative PVE by these four categories differs between leaves and roots, between microbiota and pathobiota and among the six seasonal groups.

### Identification of candidate genes associated with microbiota and pathobiota descriptors and associated enriched biological pathways

Based on a custom script (Libourel et al. 2021), we retrieved all candidate genes underlying QTLs by selecting all genes inside the QTL regions as well as the first gene upstream and the first gene downstream of these QTL regions (Data Set 11). The TAIR 10 database (http://www.arabidopsis.org/) was used as our reference. The number of candidate genes that were either specific to a single ‘plant compartment × seasonal group’ combination (single microbiota/pathobiota descriptor), or common between several ‘plant compartment × seasonal group’ combinations (several microbiota/pathobiota descriptors), were illustrated by UpSet plots using the UpSetR package in R (Conway et al. 2017).

To identify biological pathways significantly over-represented (*P* < 0.01) in each of the six ‘plant compartment × seasonal group’ combinations, each of the six lists of unique candidate genes were submitted to the classification superviewer tool on the university of Toronto website (http://bar.utoronto.ca/ntools/cgibin/ntools_classification_superviewer.cgi) using the MAPMAN classification (Provart and Zhu, 2003) (Data Set 14).

### Enrichment in signatures of local adaptation

For supporting signals of local adaptation identified by GEA analysis, we first performed for each ‘plant compartment × seasonal group’ combination, a genome-wide selection scan by estimating the XtX measure of spatial genetic differentiation among the populations. For a given SNP, XtX is a measure of the variance of the standardized population allele frequencies, which results from a rescaling based on the covariance matrix of population allele frequencies (Gautier 2015). Such rescaling allows for a robust identification of highly differentiated SNPs by correcting for the genome-wide effects of confounding demographic evolutionary forces, such as genetic drift and gene flow (Gautier 2015). For each of the 194 microbiota/pathobiota traits, we tested whether top SNPs present signatures of local adaptation by following the methodology described in (Brachi et al. 2015). More precisely, we tested whether the top SNPs were over-represented in the extreme upper tail of the XtX distribution using the formula:

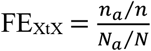

With *n* being the number of SNPs in the upper tail of the XtX distribution. In our case, we used a threshold of 0.5%. *na* is the number of top SNPs that were also in the upper tail of the XtX distribution. *N* is the total number of SNPs tested genome-wide and *Na* is the total number of top SNPs. Statistical significance of enrichment was assessed by running 10,000 null circular permutations based on the methodology described in (Hancock et al. 2011).

### Overlap with candidate genes obtained from GWAS

Based on a GWAS performed on the leaf bacterial communities of 200 Swedish *A. thaliana* accessions grown in the native habitats of four natural populations of *A. thaliana* in Sweden (Brachi et al. 2022), we retrieved a list of 880 unique candidate genes underlying 209 QTLs, by selecting all genes inside the QTL regions as well as the first gene upstream and the first gene downstream of these QTL regions.

To test whether the list of unique candidate genes obtained for each ‘plant compartment × seasonal group’ combination significantly overlaps with the list of 880 candidate genes, we first estimated the percentage of the 800 candidate genes that were in common with the list of *n* candidate genes identified in this study. To estimate the level of significance, we then created a null distribution by randomly sampling 10,000 times, *n* genes across the entire set of 27,206 genes present across the five chromosomes of *A. thaliana*.

## Acknowledgments and funding information

This project has received funding from the European Research Council (ERC) under the European Union’s Horizon 2020 research and innovation programme (grant agreement No 951444 – PATHOCOM).

## Author contributions

F.R., L.F. and C.B. planned and designed the research. F.R. performed the statistical analyses and the genome-environment analysis. F.R. wrote the manuscript, with contributions from L.F. and C.B.

