## Supplementary Information for "The genetic architecture of adaptation to leaf and root bacterial microbiota in *Arabidopsis thaliana*"

**This file includes:**

Supplementary Figures 1-5

Supplementary Tables 1-2

**Table S1.** Total number of traits related to microbiota, climate, soil physico-chemical properties and plant communities, investigated for GEA analysis.

| | | Number of traits investigted for GEA analysis | | | | |
|---|---|---|---|---|---|---|
| **Seasonal group** | **No. populations** | **microbiota/pathobiota** | **climate** | **soil** | **plant communities** | **Total** |
| fall - leaf | 73 | 42 | 6 | 14 | 35 | 97 |
| fall -root | 73 | 22 | 6 | 14 | 34 | 76 |
| spring (November ) - leaf | 69 | 40 | 6 | 14 | 35 | 95 |
| spring (November) - root | 72 | 28 | 6 | 14 | 36 | 84 |
| spring (December) - leaf | 66 | 42 | 6 | 14 | 38 | 100 |
| spring (December) - root | 66 | 20 | 6 | 14 | 38 | 78 |

**Table S2.** Differences of the percentage of variance explained (PVE) of microbiota/pathobiota traits by the demographic history of *A. thaliana*, host genetic with QTLs specific to microbiota/pathobiota traits, ecology (including climate variables, soil physico-chemical properties and descriptors of plant communities) and residuals. (A) Between the leaf and root compartment. (B) Between microbiota and pathobiota traits. (C) Between the six seasonal groups. (D) Among OTUs at the order level. (E) Among OTUs at the family level. (F) Among OTUs at the genus level. Significant *p*-values after a Bonferroni correction are indicated in red.

| **A** | leaf compartment |
|---|---|
| root compartment | 0.8517 |

| **B** | microbiota |
|---|---|
| pathobiota | 0.7409 |

| **C** | fall - leaf | fall - root | spring (November) - leaf | spring (November) - root | spring (December) - leaf |
|---|---|---|---|---|---|
| fall - root | 0.85257 | | | | |
| spring (November) - leaf | 0.77214 | 0.8881 | | | |
| spring (November) - root | 0.60547 | 0.74252 | 0.30570 | | |
| spring (December) - leaf | 0.89078 | 0.58655 | 0.76332 | 0.50662 | |
| spring (December) - root | 0.84882 | 0.70463 | 0.80064 | 0.69288 | 0.93836 |

| **D** | Burkholderiales | Pseudomonadales | Rhizobiales | Sphingomonadales |
|---|---|---|---|---|
| Pseudomonadales | 0.4802 | | | |
| Rhizobiales | 0.9149 | 0.7625 | | |
| Sphingomonadales | 0.7772 | 0.3566 | 0.6594 | |
| unclassified | 0.9998 | 0.5704 | 0.9370 | 0.6016 |

| **E** | Burkholderiales unclassified | Comamonadaceae | Oxalobacteraceae | Pseudomonadaceae | Rhizobiaceae | Runclassified |
|---|---|---|---|---|---|---|
| Comamonadaceae | 0.53826 | | | | | |
| Oxalobacteraceae | 0.89576 | 0.79813 | | | | |
| Pseudomonadaceae | 0.20224 | 0.72429 | 0.57684 | | | |
| Rhizobiaceae | 0.00063 | 0.09569 | 0.00032 | 0.00132 | | |
| Rhizobiales unclassified | 0.53437 | 0.60731 | 0.89550 | 0.74896 | 0.09769 | |
| Sphingomonadaceae | 0.71467 | 0.55395 | 0.80032 | 0.35662 | 0.08846 | 0.63480 |

| **F** | Pseudomonas | Rhizobium | Rubrivivax | Sphingomonas |
|---|---|---|---|---|
| Rhizobium | 0.00132 | | | |
| Rubrivivax | 0.49584 | **0.03817** | | |
| Sphingomonas | 0.41611 | 0.09852 | 0.29544 | |
| Variovorax | 0.11632 | 0.45732 | 0.00158 | 0.21512 |

**Supplementary Fig. 1.** Percentage of variance explained (PVE) of microbiota/pathobiota traits by the demographic history of *A. thaliana*, the individual QTLs specific to microbiota/pathobiota traits and the individual ecological variables (including climate variables, soil physico-chemical properties and descriptors of plant communities) across the six ‘plant compartment × seasonal group’ combinations. Residuals correspond to the percentage of variance unexplained by the three other categories of variables.

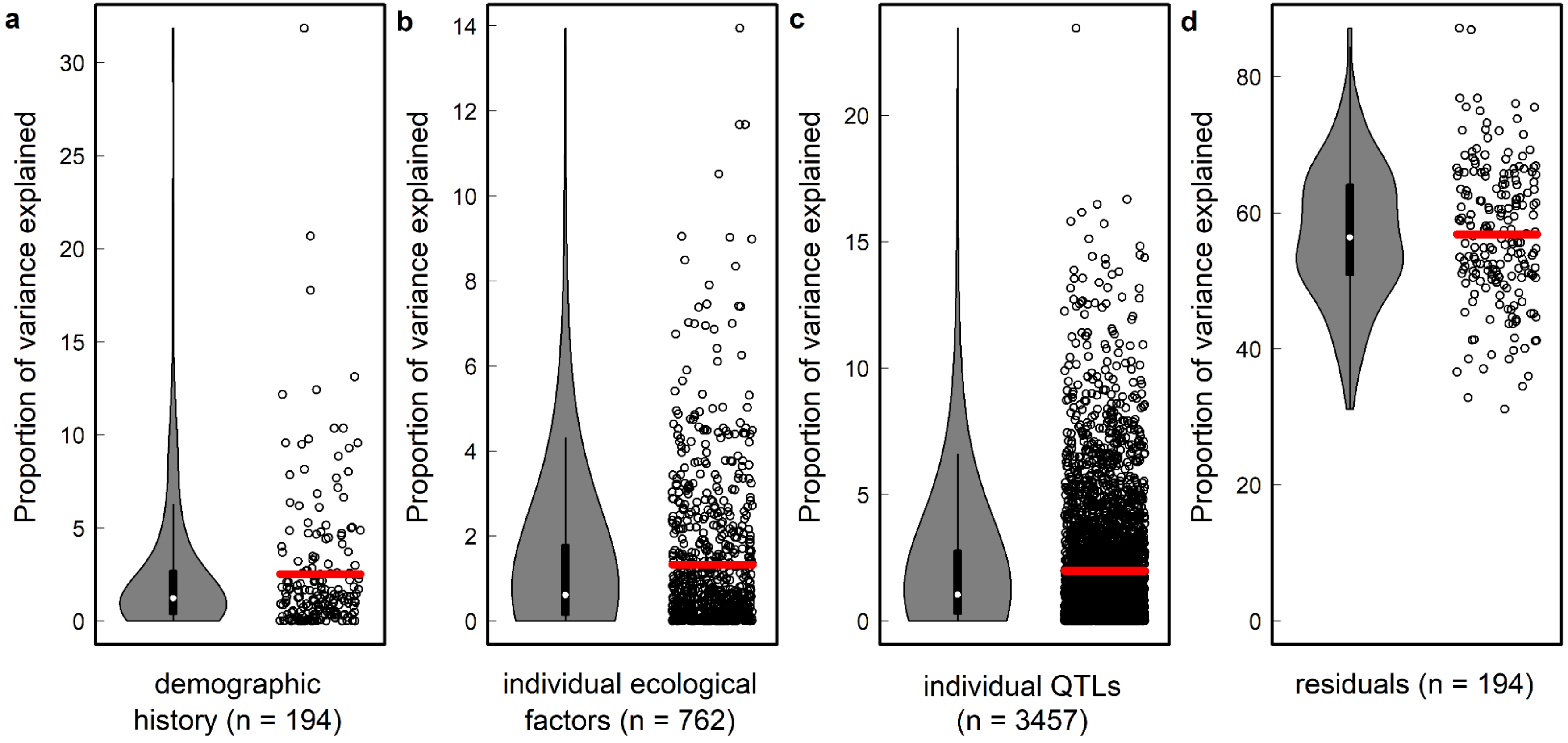

**Supplementary Fig. 2.** Cumulative percentage of variance explained (PVE) of microbiota/pathobiota traits by the demographic history of *A. thaliana*, host genetics with QTLs specific to microbiota/pathobiota traits and ecology (including climate variables, soil physico-chemical properties and descriptors of plant communities) for each 'plant compartment × seasonal group' combination. Residuals correspond to the percentage of variance unexplained by the three other categories of variables. 'FL': fall – leaf, 'FR': fall – root, 'SnovL': spring (November) – leaf, 'SnovR': spring (November) – root, 'SdecL': spring (December) – leaf, 'SdecR': spring (December) – root. For each of the four categories, the194 dots correspond to the 194 microbiota/pathobiota traits across the six 'plant compartment * seasonal group' combination.

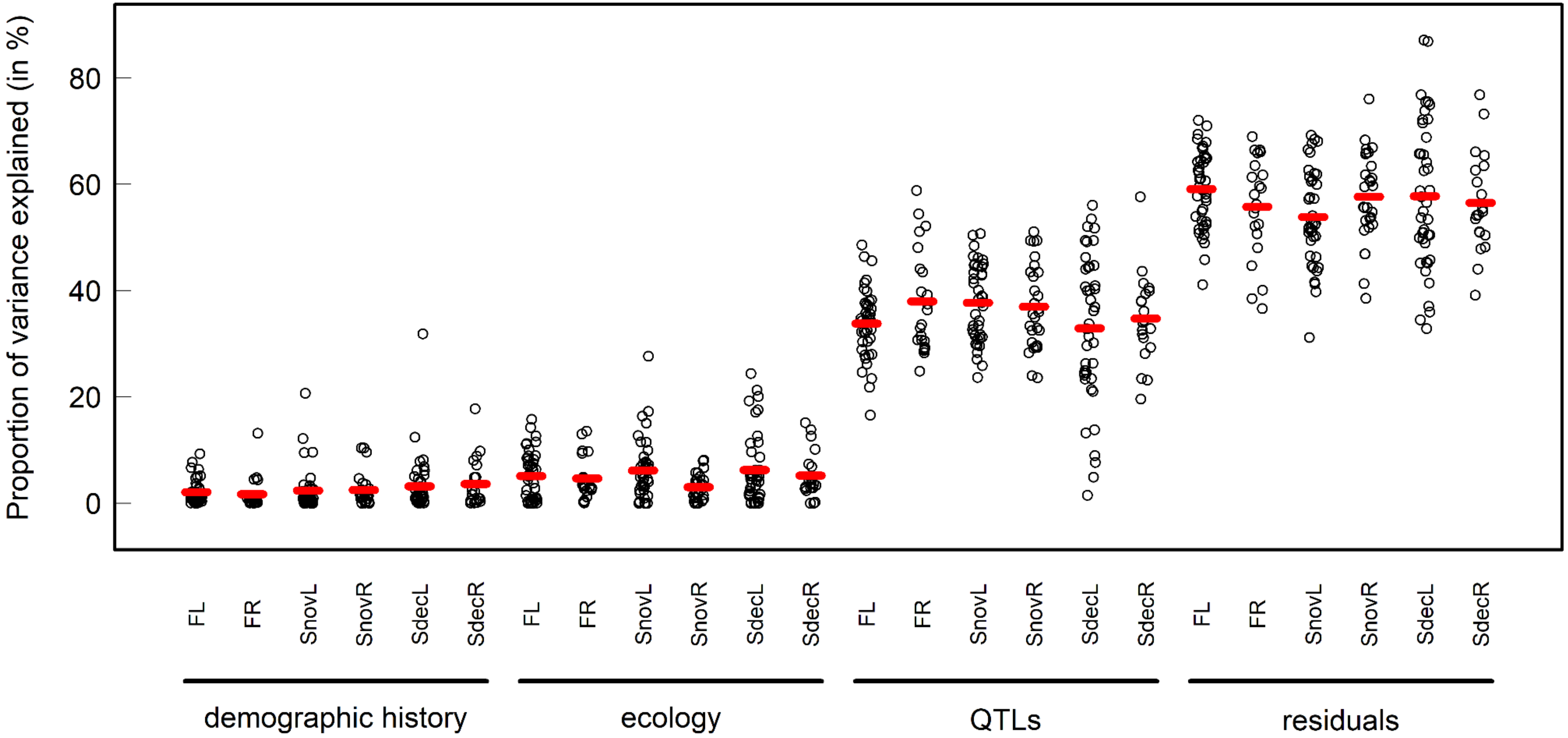

**Supplementary Fig. 3.** UpSet plots showing the intersections of the lists of candidate genes associated with microbiota/pathobiota variation, each list corresponding to one of the microbiota/pathobiota descriptors characterized on 73 populations for the seasonal group ‘fall – leaf’ (A), 73 populations for the seasonal group ‘fall – root’ (B), 69 populations for the seasonal group ‘spring (November) – leaf’ (C), 72 populations for the seasonal group ‘spring (November) – root’ (D), 66 populations for the seasonal group ‘spring (December) – leaf’ (E) and 66 populations for the seasonal group ‘spring (December) – root’ (F).

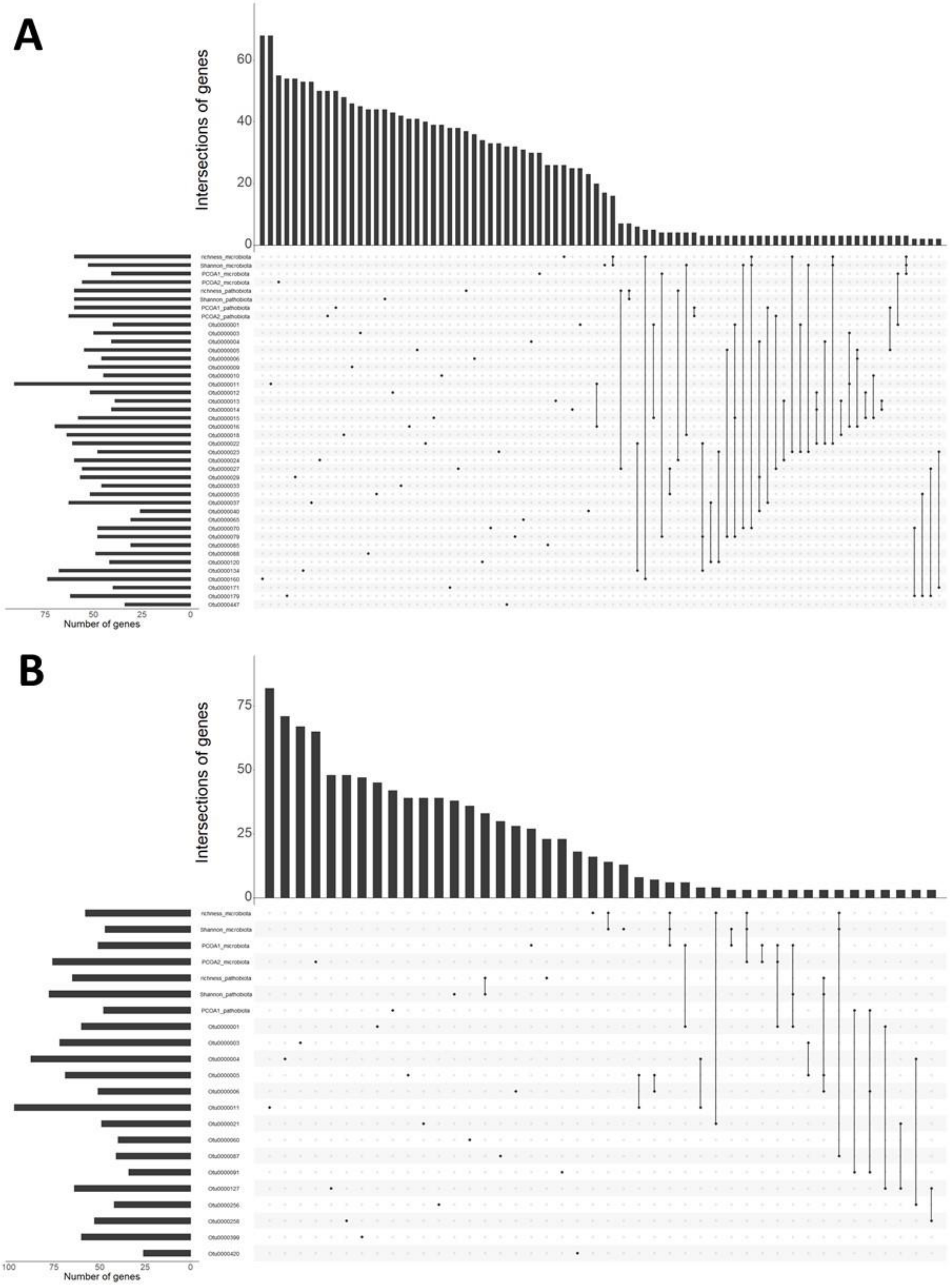

**Supplementary Fig. 3 (continued)**

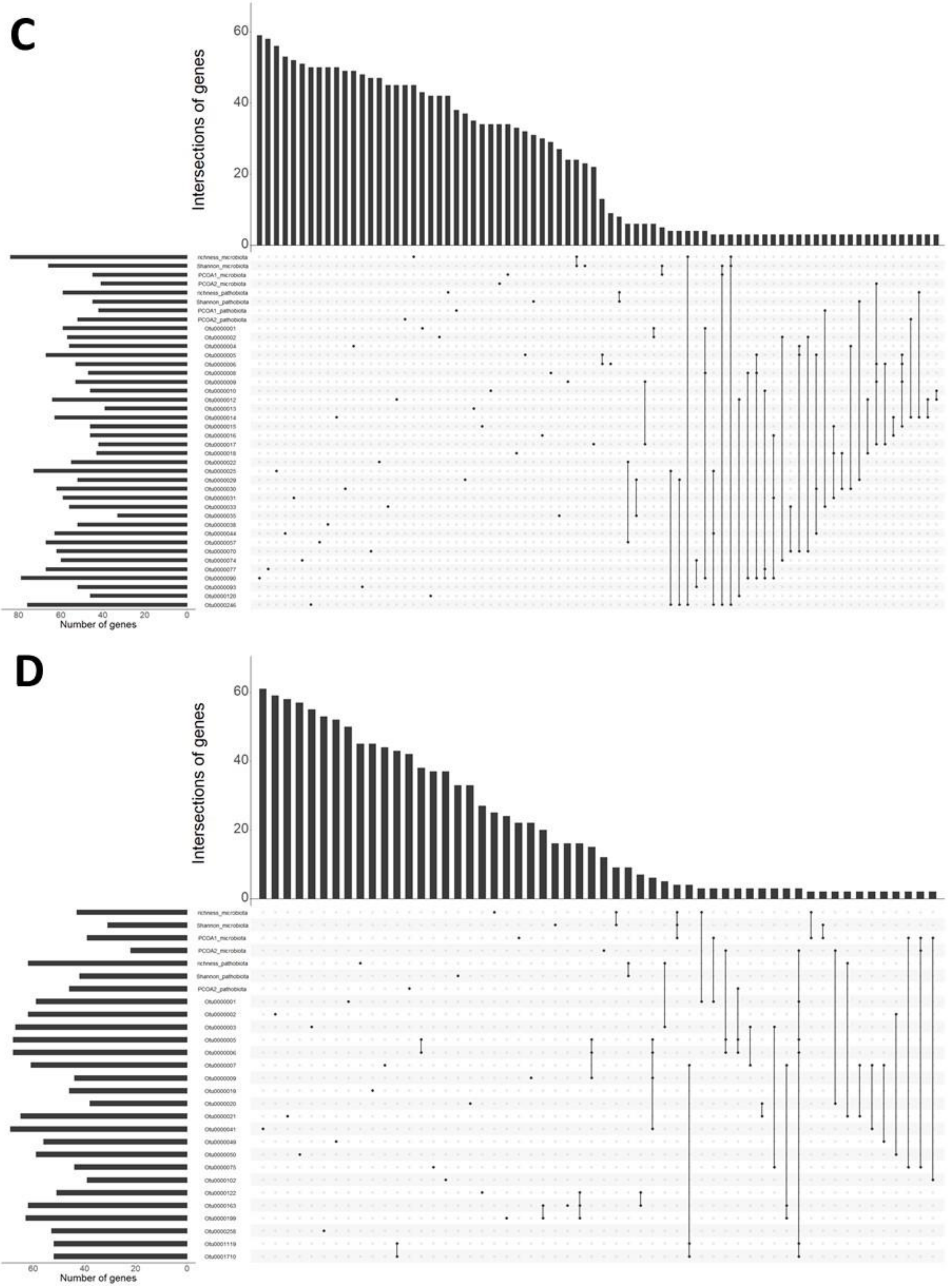

**Supplementary Fig. 3 (continued)**

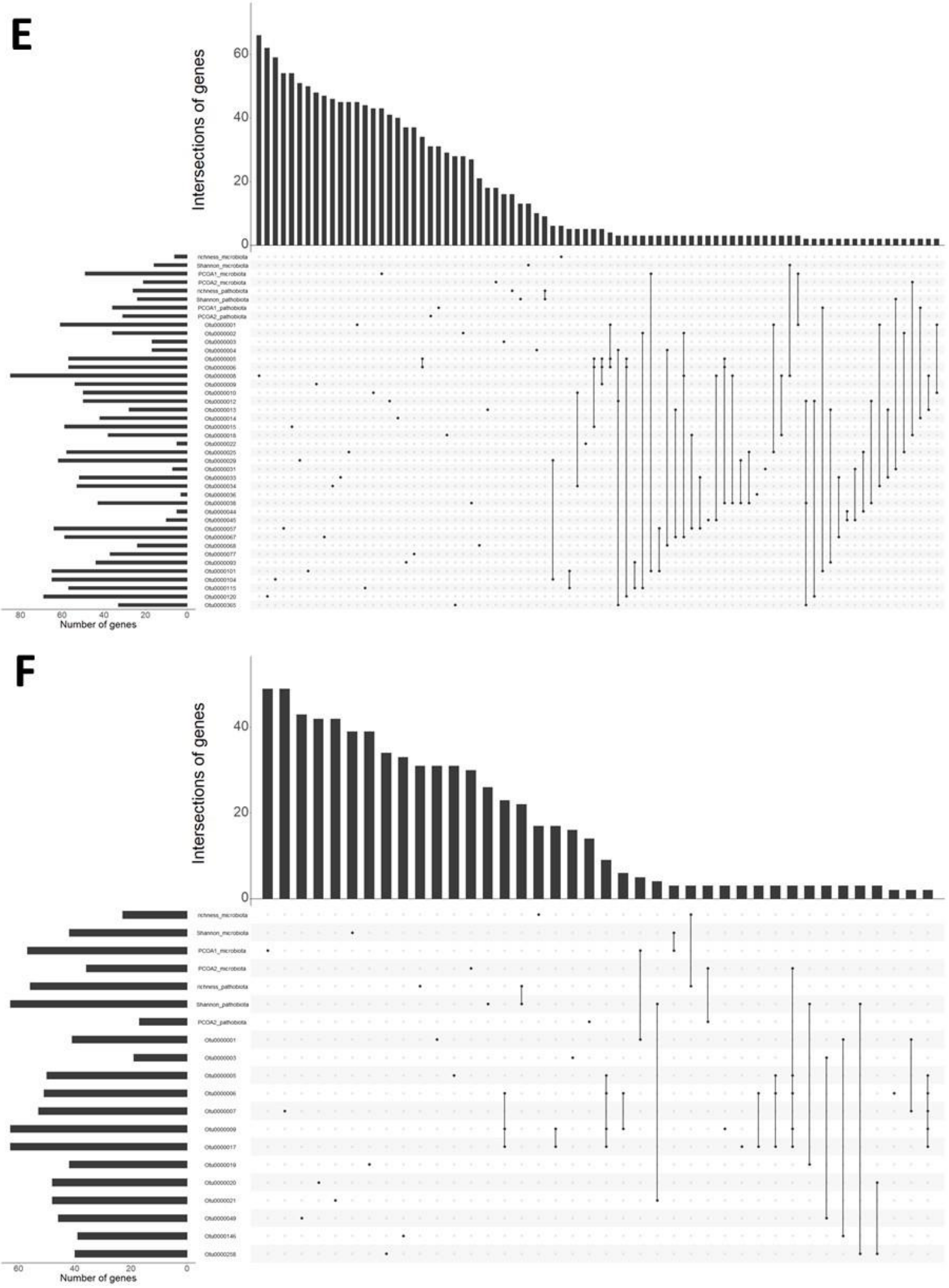

**Supplementary Fig. 4.** Manhattan plot illustrating the flexibility of genetic architecture associated with pathobiota traits in the leaf compartment in the 'spring (December)' seasonal group, i.e. Shannon index (A), composition (B) and the presence/absence of OTU8. The *x*-axis indicates along the five chromosomes, the physical position of the 1,514,789 SNPs considered for the leaf compartment in the 'spring (December)' seasonal group. The y-axis correspond to the values of the Lindley process (local score method with a tuning parameter ξ = 2). The dashed lines indicate the minimum and maximum of the five chromosome-wide significance thresholds.

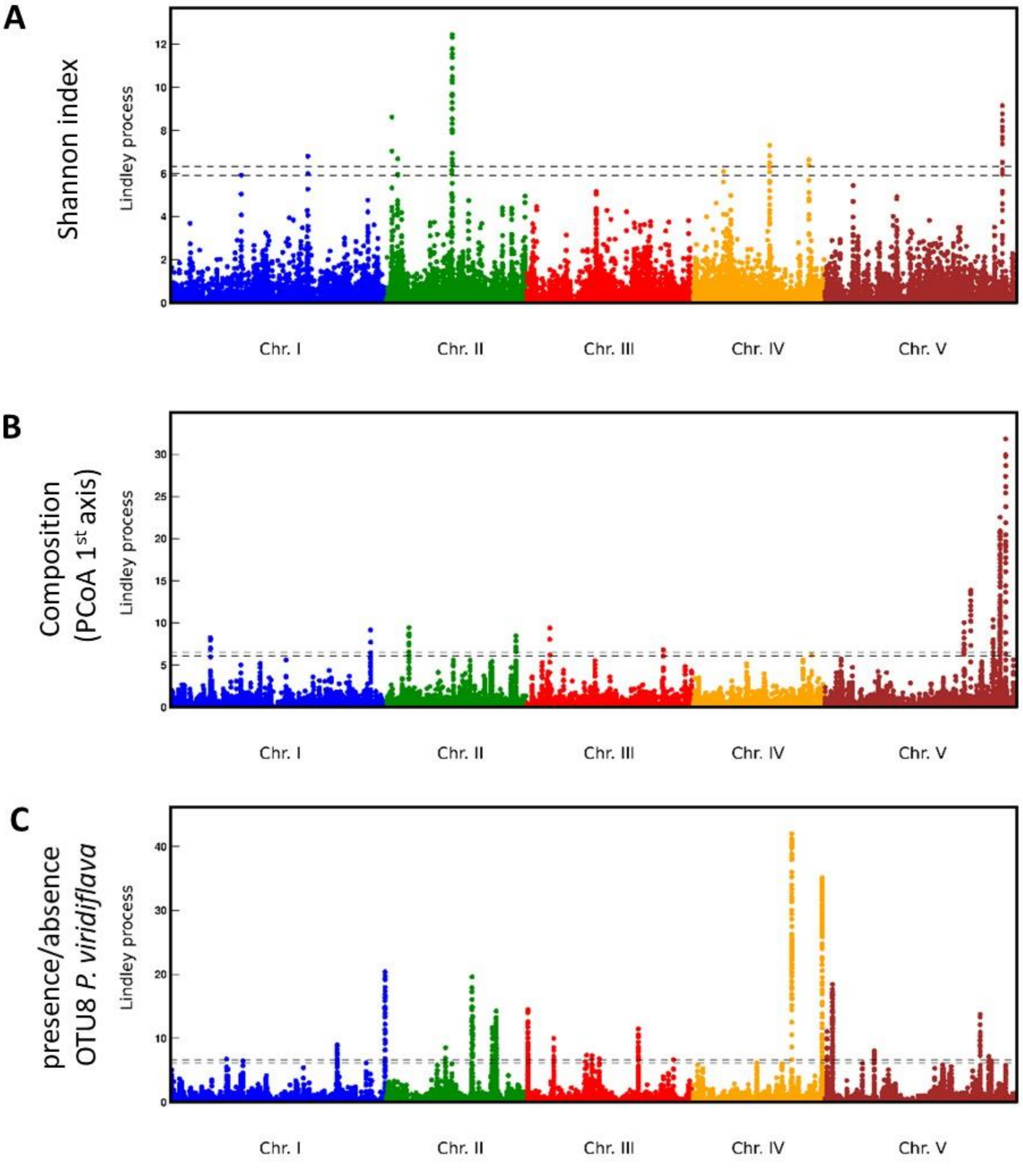

**Supplementary Fig. 5.** Cross-validation of the candidate genes identified by GEA by candidate genes identified by GWAS. (A) Fraction candidate genes obtained from a GWAS performed on the leaf bacterial communities of 200 Swedish *A. thaliana* accessions grown in the native habitats of four natural populations of *A. thaliana* in Sweden (Brachi et al. 2022) overlapping with GEA candidate genes. Large red, purple, orange and black dots correspond to the fraction of overlapped GWA candidate genes with a *p*-value $P < 0.001$, $P < 0.01$, $P < 0.05$ and $P > 0.05$, respectively. Numbers in brackets correspond to the number of unique candidate genes *n* for each category considered. Smaller gray dots represent 10,000 random sampling of *n* genes across the entire set of 27,206 genes present across the five chromosomes of *A. thaliana*. (B) Enriched biological processes for the list of unique candidate genes for each of the six 'plant compartment × seasonal group' combination, obtained with the MapMan classification SuperViewer tool. The color of the dots corresponds to the level of significance.

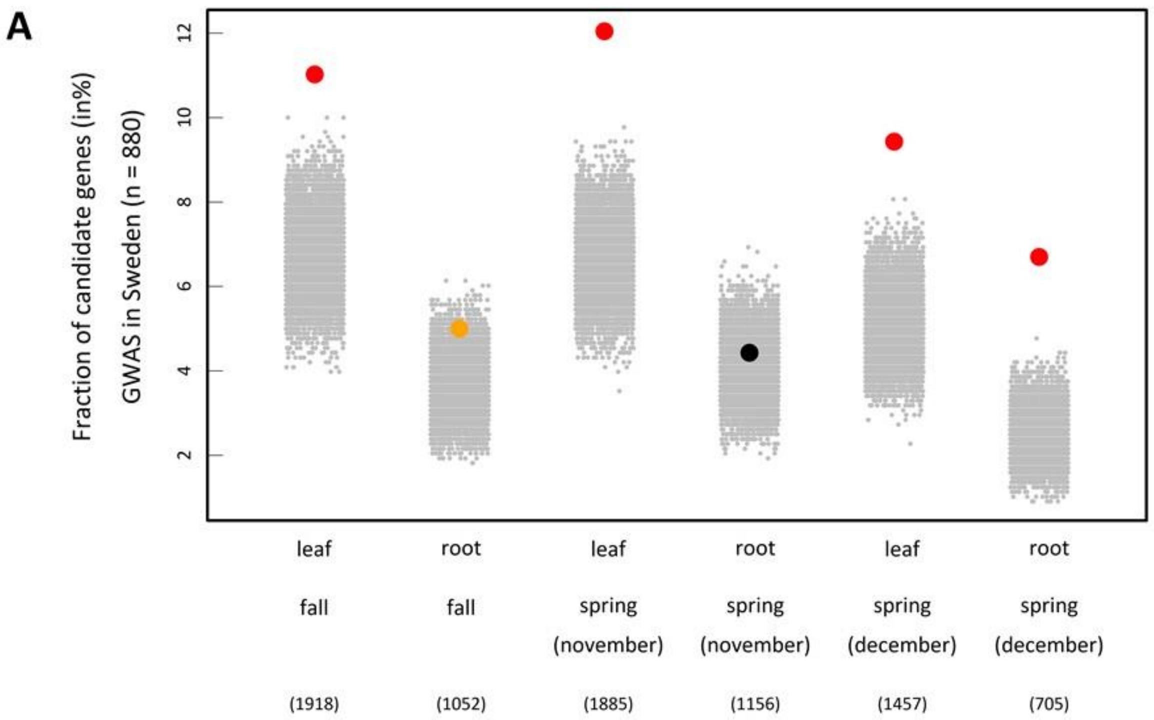


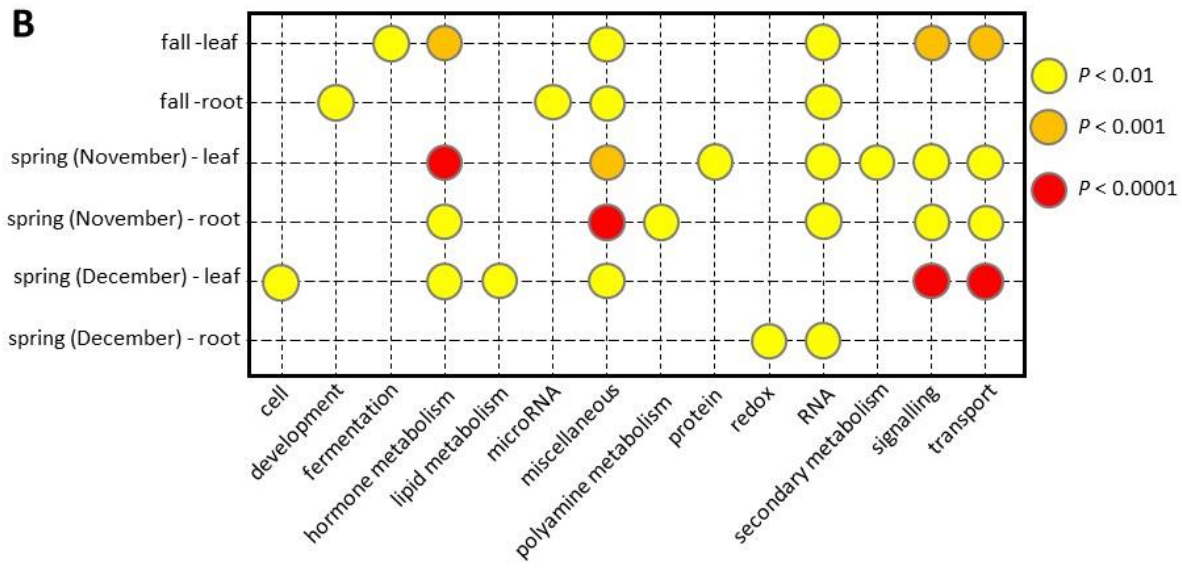
